# Structure of the lysosomal membrane fusion machinery

**DOI:** 10.1101/2022.05.05.490745

**Authors:** Dmitry Shvarev, Jannis Schoppe, Caroline König, Angela Perz, Nadia Füllbrunn, Stephan Kiontke, Lars Langemeyer, Dovile Januliene, Kilian Schnelle, Daniel Kümmel, Florian Fröhlich, Arne Moeller, Christian Ungermann

## Abstract

Lysosomes are of central importance in cellular recycling, nutrient signaling ^1,2^ and endocytosis, and are tightly connected to autophagy ^3^ and the invasion of pathogenic bacteria and viruses ^1,4^. Lysosomal fusion events are fundamental to cell survival and require HOPS, a conserved heterohexameric tethering complex ^5,6^. HOPS recognizes and binds small membrane-associated GTPases on lysosomes and organelles, and assembles membrane bound SNAREs for fusion ^7,8^. Through tethering, HOPS brings membranes in close proximity to each other and significantly increases fusion efficacy by catalysing SNARE assembly. Consequently, different HOPS mutations are causative for severe diseases ^6^. Despite its fundamental cellular duties, it remained speculative how HOPS fulfils its function as high-resolution structural data were unavailable. Here, we used cryo-electron microscopy to reveal the structure of HOPS. In the complex, two central subunits form the backbone and an assembly hub for the functional domains. Two GTPase binding units extend to opposing ends, while the SNARE binding module points to the side, resulting in a triangular shape of the complex. Unlike previously reported, HOPS is surprisingly rigid and extensive flexibility is confined to its extremities. We show that HOPS complex variants with mutations proximal to the backbone can still tether membranes but fail to efficiently promote fusion indicating, that the observed integrity of HOPS is essential to its function. In our model, the core of HOPS acts as a counter bearing between the flexible GTPase binding domains. This positions the SNARE binding module exactly between the GTPase anchored membranes to promote fusion. Our structural and functional analysis reveals the link between the spectacular architecture of HOPS and its mechanism that couples membrane tethering and SNARE assembly, to catalyse lysosomal fusion.

## Main text

Lysosomal fusion underlies a plethora of cellular processes. It is essential in the maintenance and upkeep of eukaryotic membranes and fundamental to secretion, endocytosis and autophagy. Macromolecules from different trafficking pathways end up in lysosomes where they are degraded ^2,3^. This process relies on multiple fusion events within the endomembrane system. In general, fusion depends on SNAREs, which are present on opposite membranes and zipper into four-helix bundles with the help of Sec1/Munc1 (SM) proteins ^9–11^. Prior to fusion, specialized tethering complexes establish tight links between organelles and interact with SM proteins to promote fusion ^6,8,12^. Despite their central position in trafficking, the underlying mechanics of tethering complexes and how they catalyze membrane fusion remain unresolved.

The heterohexameric HOPS complex mediates fusion of late endosomes, autophagosomes and AP-3 vesicles with mammalian lysosomes or vacuoles in yeast ^5,11,13^, and is probably the best studied tethering complex. Fusion assays using yeast vacuoles or reconstituted SNARE-bearing proteoliposomes showed that HOPS is essential for membrane fusion at physiological SNARE concentrations ^14–16^. HOPS is the target of viruses such as SARS-CoV2 ^17^, and its inactivation blocks Ebola infections ^18^. Furthermore, multiple HOPS mutations can cause severe diseases ranging from Parkinson’s to lysosomal disorders ^5,19,20^.

Five out of six HOPS subunits (Vps11, 16, 18, 39, 41) are predicted to share a similar architecture, comprising an N-terminal β-propeller and a C-terminal α-solenoid domain (Fig. 1a). Vps11 and Vps18 form the core and carry conserved C-terminal RING finger domains ^21^, which are essential for HOPS formation ^22^, but also show E3 ligase activity on their own ^23^. At the opposite sites, Vps41 and Vps39 bind to membrane anchored small GTPases (the Rab7-like Ypt7 in yeast) ^24–26^, while Vps16 and the SM protein Vps33 establish the SNARE-binding module ^7,27^. Low resolution negative-stain electron microscopy analyses revealed the overall arrangement of HOPS ^25,28^, yet were insufficient to localize the exact position of its subunits and suggested significant flexibility within the particle.

**Figure 1:**
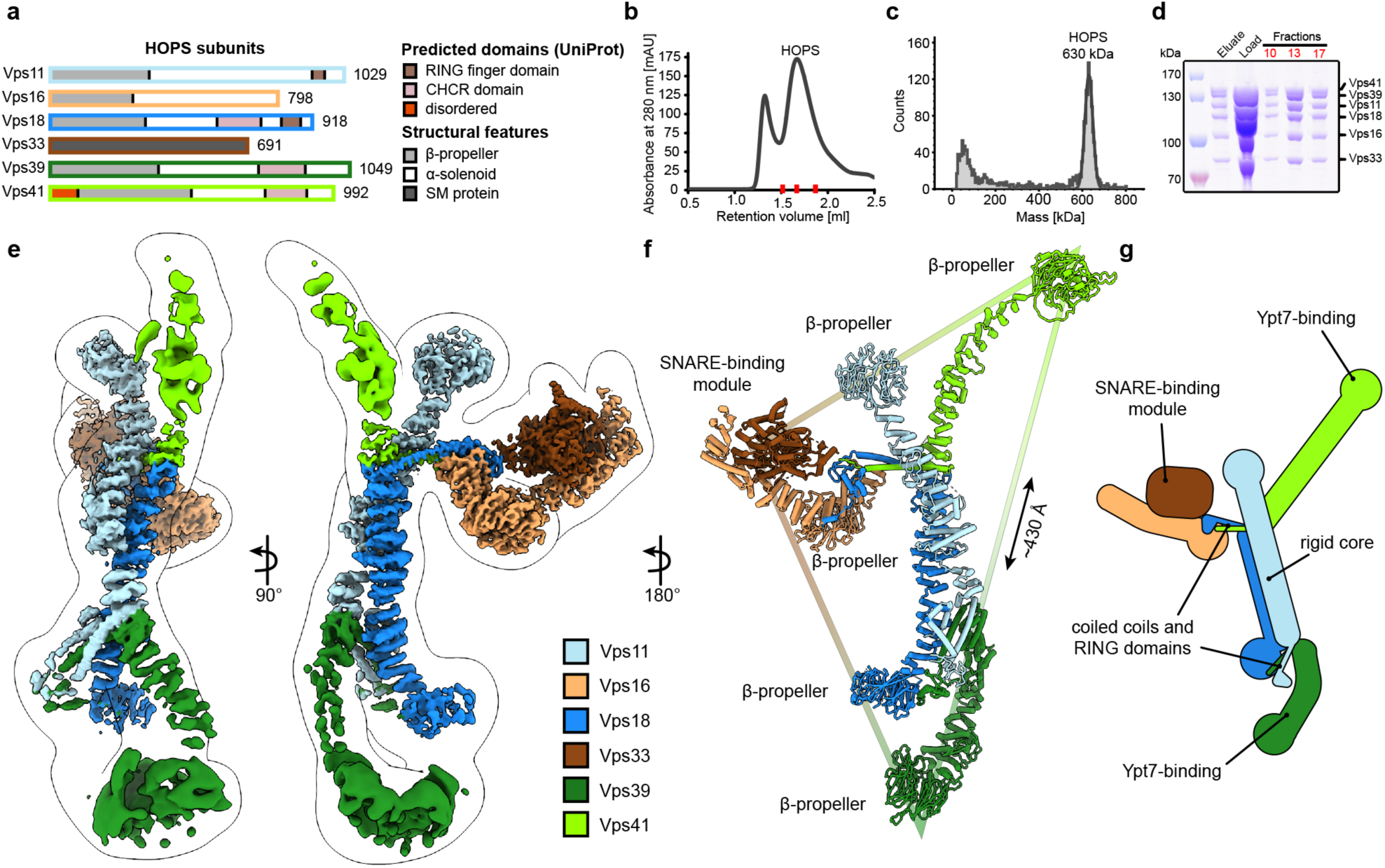
Composition and architecture of the yeast HOPS complex. **a**, Domain architecture and size of HOPS subunits. Predicted domains and structural features are indicated. **b**, Size exclusion chromatography (SEC) of the affinity-purified HOPS. Purification was done as described in Methods. **c**, Mass photometry analysis. Peak fractions from SEC were analyzed for size. **d**, Purified HOPS. Proteins from affinity purification (eluate) and SEC (red dashes in **b**) were analyzed by SDS-PAGE. **e**, Overall architecture of the HOPS complex. Composite map from local refinement maps (Extended Data Fig. 1-3) was coloured by assigned subunits. One of consensus maps used for local refinements was low-pass-filtered and is shown as a transparent envelope. **f**, Atomic model of the HOPS complex. For the N-terminal fragments of Vps41 and Vps39, which were not resolved to high resolution by local refinements, AlphaFold models are used and manually fitted into the densities of consensus maps (Extended Data Fig. 1,2). The triangular shape of the complex is highlighted with the approximate distance between the β-propellers of Vps41 and Vps39. **g**, Schematic representation of the HOPS complex indicates central features.

The way HOPS fulfils its function remains speculative, and multiple mechanisms have been proposed, including a role as a bulky membrane stressor ^15^ or, conversely, as a highly flexible membrane tether ^25,28^. In the absence of detailed structural data, it remains obscure how HOPS facilitates lysosomal fusion.

## Structure of the HOPS complex

Previously, structural studies were hampered by the low stability and flexibility of the complex, which required fixation through mild crosslinking for sample preparation and confined structural studies to negative stain analyses ^25,28^. To enable high-resolution cryo-EM of non-crosslinked HOPS, we vastly improved and accelerated our purification protocols and removed any delays during the sample preparation procedure (Fig. 1b-d). Single-particle analysis including extensive classifications followed by local refinements led to a composite structure with resolutions between 3.6 Å and 5 Å (Fig. 1e and Extended Data Fig. 1-3).

HOPS forms a largely extended, slender structure extending approximately 430 Å in height and 130 Å in width, resembling a triangular shape (Fig. 1e-g). In the center of the modular complex, Vps11 and Vps18 align antiparallel through their elongated *α*-solenoids, establishing a surprisingly rigid core (Fig. 1e-g, 2), which is in agreement with the highest resolution obtained here (Extended Data Fig. 3). Interestingly, AlphaFold predicts a long unstructured region within Vps11 (Q760 to D784), resulting in an upper and lower part of the subunit. However, this region is clearly resolved and organized in our density. In our model, the two core subunits create a central assembly hub for the four other subunits (Vps39, Vps41, Vps16, Vps33) that fulfil specific functions and localize to the periphery of the complex. The N-termini of Vps11 and Vps18 are located distally from the core and each carry a β-propeller, which can be deleted without affecting the complex formation (Fig. 3a,d, Extended Data Fig. 4). At the C-termini, both Vps11 and Vps18 have long α-helices which are followed by RING finger domains (Fig. 1e-g, 2a,b, Extended Data Fig. 5). Both features are key elements for the stability of the modular architecture and serve as anchor points for the additional subunits. In agreement, HOPS, carrying mutations in the RING finger domains of Vps11 (*vps11-1*)^29^, selectively lost Vps39, whereas only Vps41 was obtained from a similar Vps18 mutant (Extended Data Fig. 6).

**Figure 2:**
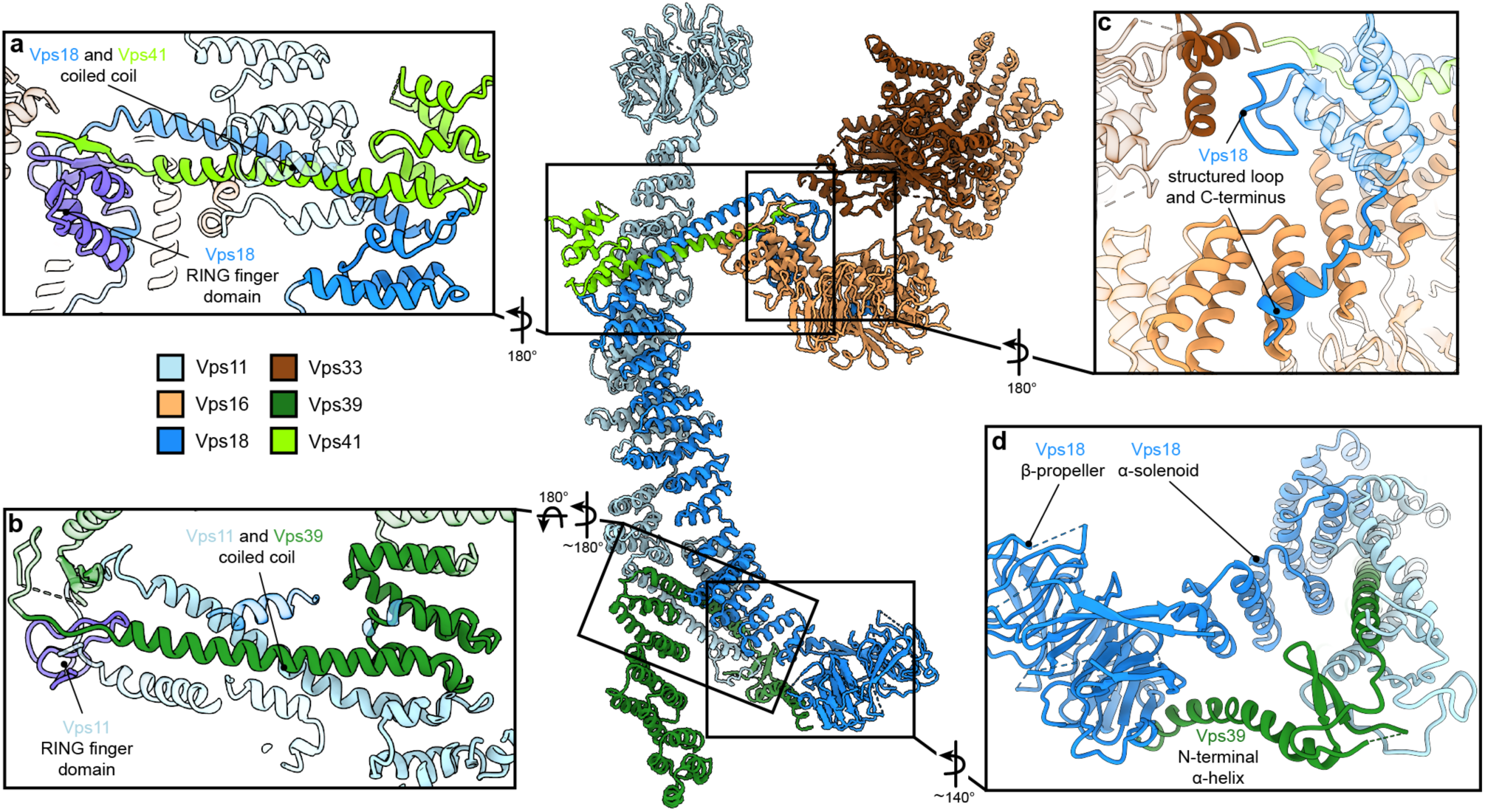
Vps11 and Vps18 C-termini as central interaction hubs for all other subunits. Atomic model of HOPS with highlighted interaction sites between subunits. **a, b**, Coiled-coil motifs followed by the RING finger domains (violet) are the key structural features of HOPS. **a**, The Vps18 C-terminal hub. Vps18 and Vps41 interact via the coiled coil and the Vps18 RING finger domain (displayed as non-transparent cartoons). **b**, The Vps11 C-terminal hub. Vps11 interacts via its RING finger domain and the coiled coil with Vps39 (displayed as non-transparent cartoons). **c**, Connection of the SNARE binding module (Vps33 and Vps16) to the backbone of HOPS via interactions with the structured loop at the RING finger domain and the C-terminus of Vps18 (displayed as non-transparent cartoons). **d**, Vps39 connects by its C-terminal helix the β-propeller of Vps18, which provides additional stability in this part of HOPS.

**Figure 3:**
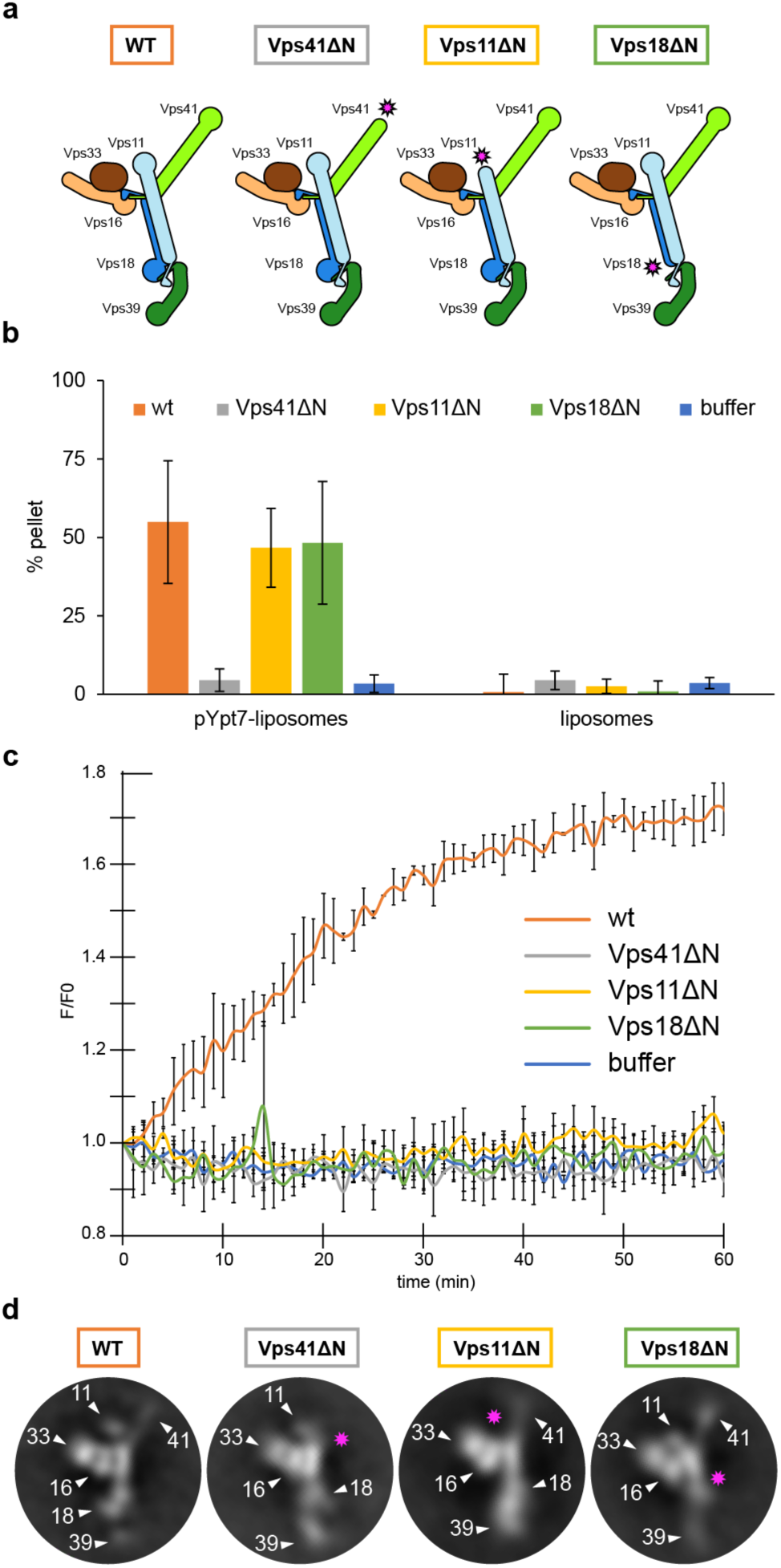
HOPS couples tethering and fusion activities. **a**, Schematic representations of HOPS wild-type (as in Fig. 1g) and mutants lacking N-terminal β-propeller domains (indicated by pink asterisks). **b**, Tethering assay. Fluorescently labeled liposomes loaded with prenylated Ypt7-GTP or none were incubated with HOPS and mutant complexes. Tethering was determined as described in Methods. **c**, Fusion assay. Fusion of proteoliposomes carrying vacuolar SNAREs were preincubated with Ypt7-GDI, GTP and Mon1-Ccz1. For fusion, HOPS wild-type or mutant and the soluble Vam7 SNARE were added 32,74. Analysis was done as described 16. See Methods. **d**, Representative 2D class averages obtained from negative-stain analyses of wild-type HOPS and mutants. Pink asterisks indicate missing densities in the mutants.

Vps41 and Vps39 provide the Ypt7-interaction sites at their peripheral N-terminal regions. Their extended C-terminal helices, similar to those of Vps11 and Vps18, are tightly interlocked through coiled-coil motifs with the long C-terminal α-helices and RING finger domains from the respective core subunits (Vps41 with Vps18, and Vps39 with Vps11) (Fig. 2a,b). Additional stability of Vps39 within the complex is provided by the interaction of the long α-helix at the C-terminus of Vps39 with the β-propeller of Vps18 (Fig. 2d). In our density, peripheral portions of both Vps41 and Vps39 are least well resolved indicating their considerable flexibility. Multi-modular classification analyses revealed angular re-orientations of about 9° for Vps41 and 20° for Vps39 (Extended Data Fig. 7a-d) relative to the core, resulting in variable positions of the N-terminal β-propellers. At the top, Vps41 reaches out by approximately 100 Å in length and, similarly, Vps39 forms an elongated arch at the bottom (Fig. 1e-g), positioning both Ypt7-interacting units at the farthest ends of the complex.

The SNARE-binding element, composed of Vps16 and the SM protein Vps33, branches out to the lateral side of the complex approximately at the center of the structure (Fig. 1e-g, 2). Vps16 shares a large interface with the coiled-coil motif formed by Vps18 and Vps41 and the N-terminus of Vps18, which is stabilized through interactions between hydrophobic and charged residues (Fig. 2c). Vps33 is in immediate contact with the structured loop of Vps18 (residues 824-831) that connects the elongated helix with the RING finger domain (Fig. 2c). This, as well as the role of RING finger domains in the interlocking of other subunits, explains, why mutations at RING domains result in devastating human diseases ^5,19,30,31^ and cause failure of correct HOPS assembly (Extended Data Fig. 6). Overall, the SNARE binding module appears to be rigidly connected to the central core, while only the short C-terminal section of Vps16 α-solenoid (residues 739-798) displays high variability and is not resolved in our structure.

## HOPS couples tethering to fusion activity

Tethering complexes bridge membranes by binding small GTPases, but also harbor or bind SM proteins ^6^. Reconstituted vacuole fusion is strictly HOPS and Ypt7-dependent at physiological SNARE concentrations ^16,32^, suggesting that HOPS is not just a tether, but part of the fusion machinery ^7,11^. However, so far it was unknown how tethering and fusion activities of HOPS are linked mechanistically. To clarify this, we analyzed the N-terminal β-propellers of Vps41 and Vps39, the likely binding sites with Ypt7 ^24,26,33^. The intrinsic low affinity between HOPS and Ypt7 ^24^ prevented reconstitution of the complex for structural studies, therefore, we instead relied on AlphaFold predictions. Additionally, we solved the structure of the β-propeller of *Chaetomium thermophilum* Vps39 by X-ray crystallography, which largely confirmed the predicted model (Extended Data Fig. 8a). Surprisingly, in the AlphaFold model, Ypt7 binding occurs at the α-solenoid of Vps39 where it does not directly interact with the β-propeller (Extended Data Fig. 8c), as originally expected. Furthermore, the binding site is placed approximately 5-6 nm above the membrane if HOPS is in an upright position ^34^. Membrane-bound Ypt7 can still reach this site due to its 10 nm long hypervariable domain (not shown in the prediction).

In the predicted complex of Vps41 (residues 1-919) with Ypt7 (residues 1-185), the GTPase binds directly to the Vps41 β-propeller, as anticipated. However, it interacts on the opposite side from the membrane-interacting amphipathic lipid-packing sensor (ALPS) motif ^35^ (Extended Data Fig. 8b), suggesting that the hypervariable region of Ypt7 is required for binding, in analogy to Vps39. Curiously, in the predicted model, the ALPS motif faces away from the membrane which would hamper membrane binding. The substantial flexibility that we observed in this region (Extended Data Fig. 7) might, therefore, be necessary to bring the ALPS motif in contact with the bilayer.

Vps39 binds Ypt7 far stronger than Vps41 ^24,36^, and may be assisted by Vps18 to sandwich Ypt7, whereas Vps41 binds to Ypt7 apparently alone (Extended Data Fig. 8d). Such a dual interaction could explain both tighter binding and a preferred orientation of HOPS on membranes ^34^. To test the functional importance, we generated HOPS complexes lacking the β-propeller of Vps11, Vps18 or Vps41 (Fig. 3a). All complexes were purified in equimolar stoichiometry (Extended Data Fig. 4a), and interacted with Ypt7, but not the Golgi Rab Ypt1 in GST-pull down assay, suggesting that at least one Rab-binding site is maintained in all truncated complexes ^16,24–26,37^ (Extended Data Fig. 4b).

To determine the activity of HOPS mutants, we compared tethering and fusion. For tethering, we incubated liposomes bearing Ypt7-GTP with each complex and quantified clustering after centrifugation ^24,34^. HOPS lacking the Vps41 β-propeller was inactive as shown ^24^, whereas HOPS with truncated Vps11 or Vps18 was fully functional (Fig. 3b). In contrast, when added to SNARE and Ypt7-GTP bearing liposomes, only wild-type HOPS, but none of the mutant complexes promoted fusion (Fig. 3c). This was particularly puzzling for HOPS lacking either the Vps18 or Vps11 β- propeller as they had full tethering activity (Fig. 3b). Therefore, we compared the structure of HOPS mutants with the wild type using negative-stain EM. Deletions of the β-propellers of Vps41, Vps11 or Vps18 indeed resulted in a loss of protein density at the expected positions, while preserving the densities of all other subunits (Fig. 3d). Interestingly, HOPS complexes lacking β-propellers in Vps11 or Vps18 showed an alteration in the relative orientation of Vps39 within the complex, which indicates a more flexible backbone, indicating that backbone integrity is essential for HOPS full activity.

Our data suggest a model of how HOPS catalyzes fusion at lysosomes and vacuoles (Fig. 4). HOPS three major ligand binding sites are arranged in a triangular fashion. While the two Ypt7-binding sites show significant conformational variability, the SNARE-interacting unit is rigidly connected to the stiff backbone. For tethering, HOPS Vps39 and Vps41 bind Ypt7 on target membranes. During this process, HOPS remains upright on membranes ^34^. Then, the SM protein Vps33 and possibly other sites on HOPS ^7,24,38,39^ bind SNAREs from the opposing membranes, and zipper them up toward their membrane anchor (Fig. 4). Note, that this process can be blocked by Orf3a in the COVID-19 SARS-CoV-2 virus ^17^. During zippering, the stiff HOPS backbone dampens the movement of the vesicles and acts as a lever arm holding on to SNAREs ^15^. This three-point arrangement results in membrane stress, and can explain how HOPS catalyzes membrane fusion (Fig. 4). The physiological function of HOPS can be bypassed if large complexes are redirected to SNAREs at the fusion site ^15,40^, which can even promote fusion of deficient SNARE complexes ^41,42^. Zippered SNAREs may then dissociate from HOPS and allow access for α-SNAP and NSF to recycle SNAREs ^43^.

**Figure 4:**
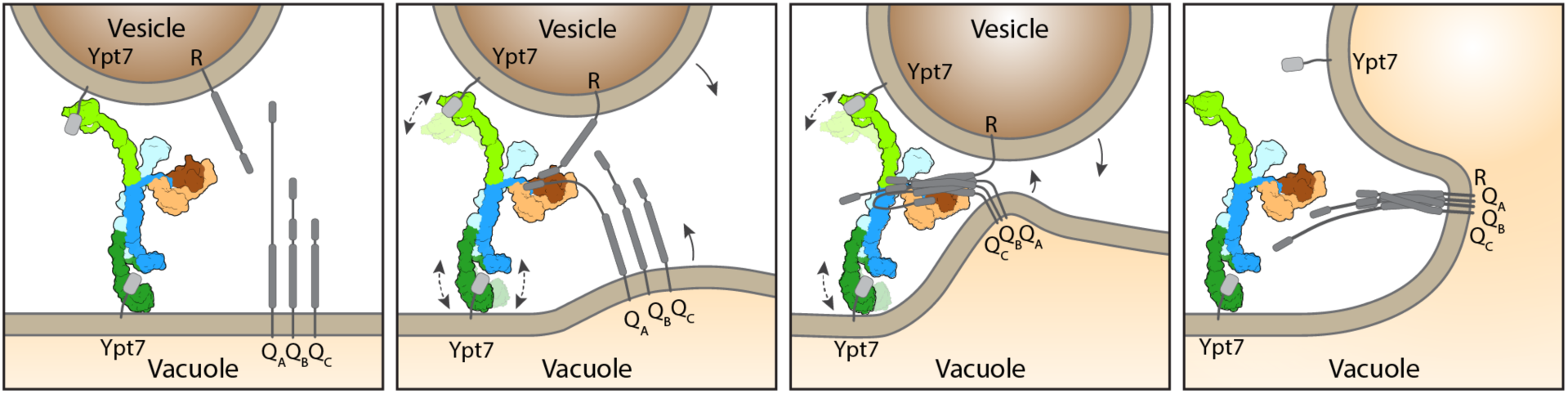
Model for HOPS-mediated membrane tethering and fusion. The HOPS complex binds to Ypt7 on the vacuole and vesicles via Vps39 (dark green) and Vps41 (light green). SNARE proteins are recruited by the SNARE-binding module (dark and light brown). The rigid core of HOPS keeps the membranes in place, while Vps41 and Vps39 function as dampers due to their limited flexibility. Consequentially, zippering of SNAREs on Vps33 (dark brown) occurs against the lever of the rigid HOPS backbone, causes membrane stress and thus catalyzes fusion. HOPS may let go of membranes thereafter.

## Conclusion

How tethering complexes catalyze fusion poses a long-standing question in the field. The process necessitates binding of two opposing membranes and exact coordination of the zippering procedure of membrane bound SNAREs. Previous analyses suggested strong flexibility along the entire structure, which was interpreted as a hallmark of tethering complexes and an essential prerequisite to their function ^25,28,34,44^. Structural models such as the “spaghetti dancer” ^6^ supported the importance of HOPS structural flexibility. Instead, our data shows that HOPS flexibility is limited, and its backbone and the SNARE-binding module are stiff, while the membrane interacting units dampen motion and stabilize the SNARE interaction, which appears to be necessary for fusion. Mammalian HOPS contains the same six subunits as yeast HOPS, which are generally highly conserved and will therefore function similarly ^5,45^. Our structure can hence be used to map disease prone mutations (Extended Data Fig. 9) and may thus explain the consequences on HOPS function.

Some tethering complexes such as CORVET, CHEVI or FERARI have attached SM proteins ^5,46^, others such as COG, GARP, Exocyst and Dsl cooperate with SM proteins ^6^, which may catalyze fusion similar to HOPS. Moreover, the overarching principle shown here for HOPS is not limited to lysosomal fusion but extends to synaptic transmission, where the tether Munc13 and the SM protein Munc18 cooperatively catalyze SNARE-mediated fusion ^47–50^. Therefore, our insights provide an exciting blueprint to understand HOPS function, dynamics and regulation both in fusion and in other functions of its subunits ^51–54^, and imply a general role of tethering complexes as a catalytic part of the fusion machinery.

## Methods

### Yeast strains

Yeast strains used in this study are listed in Table S1. In general, HOPS subunits were expressed under the control of the GAL1 promoter according to the standard protocol ^55^. For HOPS subunit truncation (Vps41, Vps11 and Vps18) of the N-terminal part, the GAL1 promotor was inserted into the genome at the respective position. The 3x-FLAG Tag was attached to the HOPS subunit Vps41, except for the 18 ΔN mutant.

### Protein expression and purification from *Escherichia coli*

Rab GTPases for pulldown or tethering and fusion assays were expressed in *E. coli* BL21 (DE3) Rosetta cells. Cells were grown in Luria broth (LB) medium complemented with 35 μg/ml kanamycin and 30 μg/ml chloramphenicol. Cultures were induced at OD_600_ = 0.6 with 0.5 mM isopropyl-β-d-thiogalactoside (IPTG) overnight at 16°C before harvesting by centrifugation (4,800 *g*, 10 min, 4 °C). Cells resuspended in buffer (150 mM NaCl, 50 mM HEPES/NaOH, pH 7.4, 10 % glycerol, 1 mM PMSF and 0.5-fold protease inhibitor mixture (PIC)) were lysed by a Microfluidizer (Microfluidics Inc.) and centrifugated at 25,000 *g*, 30 min, 4 °C. Supernatants were incubated with glutathione Sepharose (GSH) fast flow beads (GE-Healthcare) for GST-tagged proteins or nickel– nitriloacetic acid (Ni-NTA) agarose (Qiagen) for His-tagged proteins for 2 h at 4°C. The proteins were eluted with buffer (150 mM NaCl, 50 mM HEPES/NaOH, pH 7.4, 10 % glycerol) containing 25 mM glutathione or 300 mM imidazole. Buffer was exchanged via a PD10 column (GE Healthcare). For tag cleavage, TEV protease was added after washing and incubated over night. All proteins were stored at -80 °C.

### Purification of the 3xFLAG-tagged HOPS complex variants from yeast

Two liters of yeast peptone (YP) medium containing 2 % galactose (v/v) were inoculated with 6 ml of an overnight culture. Cells were grown for 24 h and harvested by centrifugation (4,800 *g*, 10 min, 4 °C). Pellets were washed with cold HOPS purification buffer (HPB, 1 M NaCl, 20 mM HEPES/NaOH, pH 7.4, 1.5 mM MgCl_2_, 5% (v/v) glycerol). The pellet was resuspended in a 1:1 ratio (w/v) in HPB supplemented with 1 mM phenylmethylsulfonylfluoride (PMSF), 1× FY protease inhibitor mix (Serva) and 1 mM dithiothreitol (AppliChem GmbH) and afterwards dropwise frozen in liquid nitrogen before being lysed in a freezer mill cooled with liquid nitrogen (SPEX SamplePrep LLC). The powder was thawed on ice and resuspended in HPB supplemented with 1 mM PMSF, 1× FY and 1 mM DTT using a glass pipette, followed by two centrifugation steps at 5,000 and 15,000 *g* at 4 °C for 10 and 20 min, respectively. After centrifugation, the supernatant was added to 2 ml of anti-FLAG M2 affinity gel (Sigma-Aldrich) and gently agitated for 45 min at 4 °C on a nutator. Beads were briefly centrifuged (500 *g*, 1 min, 4 °C) and the supernatant was removed. Beads were transferred to a 2.5 ml MoBiCol column (MoBiTec) and washed with 25 ml of HOPS washing buffer (HWB, 1 M NaCl, 20 mM HEPES/NaOH, pH 7.4, 1.5 mM MgCl_2_, 20% (v/v) glycerol). FLAG-peptide was added and incubated on a turning wheel for 40 min at 4 °C. The eluate was collected by centrifugation (150 *g*, 30 s, 4 °C) and concentrated in a Vivaspin 100 kDa MWCO concentrator (Satorius), which was previously incubated for 45 min with HWB containing 1% TX-100. The concentrated eluate was applied to a Superose 6 Increase 15/150 column (Cytiva) for size exclusion chromatography (SEC) and eluted in 50 μl fractions using ÄKTA go purification system (Cytiva). Peak fractions were used for further analysis.

### Mass photometry

Mass photometry experiments were done using a Refeyn TwoMP (Refeyn Ltd.). Data was obtained using AcquireMP software and analysed using DiscoverMP (both Refeyn Ltd.). Glass coverslips were used for sample analysis. Perforated silicone gaskets were placed on the coverslips to form wells for every sample to be measured. Samples were evaluated at a final concentration of 10 nM in a total volume of 20 μl in the buffer used for SEC.

### Cryo-EM sample preparation and data acquisition

Prior to cryo-EM analysis, HOPS samples were tested by negative-stain electron microscopy, using 2% (w/v) uranyl formate solution as previously described ^56^. Negative-stain micrographs were recorded manually on a JEM-2100Plus transmission electron microscope (JEOL), operating at 200 kV and equipped with a XAROSA CMOS 20 Megapixel Camera (EMSIS) at a nominal magnification of 30,000 (3.12 Å per pixel). For cryo-EM, 3 μl of 0.6–0.9 mg/ml of freshly purified wt HOPS complex were applied onto glow-discharged CF grids (R1.2/1.3) (EMS) and immediately plunge-frozen in liquid ethane using a Vitrobot Mark IV (Thermo Fisher Scientific) with the environmental chamber set to 100% humidity and 4°C. Micrographs were recorded automatically with EPU (Thermo Fisher Scientific), using a Glacios microscope (Thermo Fisher Scientific) operated at 200 kV and equipped with a Selectris energy filter and a Falcon 4 detector (both Thermo Fisher Scientific). Images were recorded in Electron Event Representation (EER) mode at a nominal magnification of 130,000 (0.924 Å per pixel) in the defocus range from −0.8 to −2.8 μm with an exposure time of 8.30 s resulting in a total electron dose of approximately 50 e^−^ Å^−2^.

### Cryo-EM image processing

All cryo-EM data processing (Extended Data Fig. 1) was performed in cryoSPARC v3.3.1^57^. For all collected movies, patch motion correction (EER upsampling factor 1, EER number of fractions 40) and patch contrast transfer function (CTF) estimation were performed using cryoSPARC implementations. To solve the structure of the core part of HOPS (Extended Data Fig. 1e,f, 2), reference-free blob particle picking on 2186 pre-processed movies of the first dataset and particle extraction using a box size of 672 pixels (px, 0.924 Å per pixel) binned to 128 px was performed. Extracted particles were subjected to 2D classification to eliminate bad picks. Selected good 2D classes (representative good 2D classes are shown in Extended Data Fig. 3a) were used for template particle picking on 2186 movies from the first dataset combined with additional 6580 movies from the second dataset, preprocessed alike. After removal of duplicates, picked particles were extracted with the same box size and subjected to rounds of 2D classification, ab-initio reconstruction with multiple classes and 3D heterogeneous refinement to remove false positive particle picks. From the best class, particles were extracted using a box size of 672 px (0.924 Å per pixel) binned to 320 px and subjected to 2D classification. Particles from this 2D classification were also used for flexibility analysis (see below). 407,996 selected particles from good 2D classes were used for ab-initio reconstruction with six classes and followed by 3D heterogeneous refinement. This heterogeneous refinement resulted in two best classes containing 130,009 and 116,151 particles, respectively, which were further refined individually. For each of both classes, a Non-Uniform (NU)-refinement was performed, followed by particle extraction with a box size of 672 px (0.924 Å per pixel, without binning) with homogeneous and NU-refinement afterwards. Resulting consensus maps were used for local refinements of different parts of the structure (Extended Data Fig. 1a, 2, 3b-e). First consensus NU-refinement that reached global resolution of 4.2 Å (Fourier shell correlation (FSC) = 0.143) was used for local refinements of the upper “core” part of the complex (4 Å) and the SNARE-binding module (3.6 Å), the second consensus NU-refinement, resolved to 4.4 Å (FSC = 0.143), was used for local refinements resolving bottom parts of the complex (4.4 and 5 Å).

To better resolve distal parts of the complex, flexible Vps39 and Vps41 N-terminal fragments, the following approach was used (Extended Data Fig. 1b). Here, micrographs from first two datasets were combined with micrographs from two additional datasets of 2841 and 8338 movies. After template picking, about 3,896 million particles were extracted using a box size of 882 pixels (0.924 Å per pixel) and used for rounds of heterogeneous, homogeneous and NU-refinements to obtain 3D reconstructions, which best resolve Vps39 and Vps41. At these steps, binning was applied for particle extractions. Then, particles belonging to one of such best classes were subjected to a round of NU-refinement followed by 3D variability analyses using masks covering either Vps39 or Vps41. The further 3D variability display procedures were used to better sort particles. Finally, the best particles were extracted with the same box size (882 pixels, 0.924 Å per pixel) without binning and used for local refinements of Vps39 or Vps41 (Extended Data Fig. 1b, 3f,g).

For all local refinements, masks were generated in UCSF Chimera ^58^. During processing, no symmetry was applied. The quality of final maps is demonstrated in Extended Data Fig. 3. All FSC curves were generated in cryoSPARC. Local resolutions of locally-refined maps (Extended Data Fig. 3b-g) were estimated in cryoSPARC and analyzed in UCSF ChimeraX ^59^. Dataset statistics can be found in Table S2.

To analyze the flexibility of the complex, a different processing scheme was applied (Extended Data Fig. 1a, dashed arrows). For this, a set of 383,881 good particles was selected after 2D classification of 320 px-binned particles (initial box size of 672 px with 0.924 Å per pixel). These particles were subjected to either ab-initio reconstruction and heterogeneous refinement with ten classes (Extended Data Fig. 7a,b) or several refinement rounds followed by 3D variability analysis (Extended Data Fig. 7d). Prior to 3D variability analysis, ab-initio reconstruction with one class, homogeneous and NU-refinement were performed.

### Model building and refinement

Models of HOPS subunits were initially generated using AlphaFold ^60,61^ and docked into locally-refined maps using “Fit in Map” tool in UCSF ChimeraX ^59^. The N-terminal parts of Vps41 (residues 1-863) and Vps39 (residues 1-700) and the C-terminal part of Vps16 (from residue 739) with no well-resolved densities assigned, were removed. The AlphaFold model of Vps11 was initially split in two parts (“Vps11top”, residues 1-760, and “Vps11bottom”, residues 784-1025), which were first refined separately; the region predicted by AlphaFold as unfolded (residues 761-783) was deleted. Fitting of the C-terminal parts of Vps11 (residues 784-1025) and Vps39 (residues 701-1045) was improved using Namdinator ^62^. Afterwards, models of single proteins were manually adjusted and refined in COOT ^63^, followed by iterative rounds of refinements against corresponding locally-refined and their composite maps in Phenix ^64^ and COOT. In Phenix Graphical User Interface, real space refine tool ^65^ with or without option “rigid body” was used. After several refinement rounds, two separate models were created by joining models of the upper (Vps33, Vps16, Vps41, Vps18, Vps11top) and lower (Vps11bottom, Vps39) parts of the complex. These partly-combined models were subjected to further several iterations of refinements in Phenix and COOT. Afterwards, the two refined models were fused into a single model, which was again refined in Phenix and COOT. Sequences of the model in the bottom part of the complex were changed to polyalanines (residues 784-1025 in Vps11, residues 1-493 in Vps18, and residues 701-1045 in Vps39), since no assignment of side chains was possible at the resolution obtained there. In other parts of the complex, where blurred densities did not allow unambiguous model building, respective short fragments of the model were also replaced by polyalanine chains or deleted. Afterwards, the initially deleted part of Vps11 (residues 761-783) was built *de novo* according to the cryo-EM density (residues 769-783 were replaced by alanines). Finally, the complete model was subjected to another round of refinement in Phenix followed by manual refining in COOT. Model validation was performed using MolProbity ^66^. Figures were prepared using UCSF ChimeraX. Model refinement and validation statistics can be found in Table S2.

### ALFA Pulldowns for mass spectrometry

One liter of yeast peptone (YP) medium containing 2 % glucose (v/v) was inoculated with an overnight preculture. Cells were grown to OD_600_ 1 at 26 °C, followed by 1 h incubation at 38 °C. Cultures were harvested by centrifugation at 4,800 *g* for 10 min at 4 °C). Pellets were washed with cold Pulldown buffer (PB, 150 mM KAc, 20 mM HEPES/NaOH, pH 7.4, 5% (v/v) glycerol, 25 mM CHAPS). The pellet was resuspended in a 1:1 ratio (w/v) in PB supplemented with Complete Protease Inhibitor Cocktail (Roche) and afterwards dropwise frozen in liquid nitrogen before lysed in 6875D LARGE FREEZER/MILL® (SPEX SamplePrep LLC). Powder was thawed on ice and resuspended in PB by using a glass pipette followed by two centrifugation steps at 5,000 and 15,000 *g* at 4 °C for 10 and 20 min. The supernatant was added to 12.5 μl prewashed ALFA Selector ST beads (2,500 *g*, 2 min, 4 °C) (NanoTag Biotechnologies) and incubated for 15 min at 4 °C while rotating on a turning wheel. After incubation, beads were washed twice in PB and four times in PB without CHAPS. Samples were digested using PreOmics sample kit (iST Kit, preomics) and analysed in Q ExativePlus mass spectrometer (Thermo).

### GST Pulldowns

Nucleotide specific interaction of the Rab-GTPase Ypt7 with purified HOPS variants was analysed in GST pulldowns using GST-Ypt7 or GST-Ypt1 as a negative control. 125 μg purified Rab-GTPases were preloaded with 1 mM GDP or GTP in the presence of 20 mM EDTA and Wash buffer (150 mM NaCl, 50 mM HEPES-NaOH, pH 7.4, 2 mM MgCl_2_, 0.1 % Triton X-100) in a water bath for 30 min at 30 °C. For nucleotide stabilization, 25 mM MgCl_2_ was added. Prewashed GSH Sepharose 4B (GE Healthcare) were added to loaded Rab-GTPases and incubated for 1 h at 4 °C on a turning wheel. Beads were centrifuged for 1 min at 300 *g* before adding 25 μg of respective HOPS variants, followed by 1.5 h incubation at 4 °C on a turning wheel. Beads were washed three times with Wash buffer, followed by two elution steps with 600 μl Wash buffer containing 20 mM EDTA. After an incubation at room temperature for 20 min while rotating, the supernatant was TCA precipitated. 10 % of the final sample was loaded on a 7.5 % SDS gel next to 5 % of protein input. Samples were analysed via western blot using antibodies against the FLAG-Tag. Beads were boiled in 50 μl Laemmli buffer. 2 % of the sample was loaded on a 11 % SDS gel for Coomassie staining as Rab-GTPase loading control. Bands were quantified relative to the Rab-GTPase content.

### Tethering assay

HOPS mediated tethering assays were performed as described ^34^. For this, ATTO488 labelled liposomes were prepared and loaded with prenylated Ypt7 ^32^. 50 nmole liposomes were incubated with 50 pmole pYpt7:GDI complex together with GTP for 30 min at 27 °C. For reactions, 0.17 mM Ypt7-loaded liposomes were incubated with different concentrations of HOPS complex or buffer (300 mM NaCl, 20 mM HEPES-NaOH, pH 7.4, 1.5 mM MgCl_2_, 10 % (v/v) glycerol) for 10 min at 27 °C, followed by sedimentation for 5 min at 1,000 *g*. The overall tethered liposomes in the pellet fraction were determined in a SpectraMax M3 fluorescence plate reader (Molecular devices) comparing the ATTO488 fluorescent signal of the supernatant before and after sedimentation.

### Fusion assays

Fusion assays and the purification of all proteins was performed as described ^32^ with a protein to lipid ratio of 1:8,000. Reconstituted proteoliposomes (RPLs) were composed of the vacuole mimicking lipid mix (VML) ^67^. One population of RPLs carried the SNARE Nyv1 and the other set contained Vti1 and Vam3. RPLs were preloaded with prenylated Ypt7 with the help of 100 mM Mon1-Ccz1 and 0.5 mM GTP ^32^. Then, 25 nM HOPS complex, 50 nM Sec18, 600 nM Sec17, and finally 100 nM Vam7 were added. Fusion of liposomes was followed by content mixing of the RPLs and subsequent increase in fluorescence which was monitored in a SpectraMax M3 fluorescence plate reader (Molecular devices).

### Cloning and protein purification of CtVPS391-500

Codon-optimized synthetic DNA (GenScript) encoding amino acids 1-500 of Vps39 from *Chaetomium thermophilum* (CtVp391-500; NCBI XP_006691033) was subcloned into a modified pET28a expression vector yielding an N-terminally His6-SUMO-tagged fusion protein (His6-SUMO-CtVps391-500). In short, His6-SUMO-CtVps39 was purified by Ni-NTA affinity chromatography followed by proteolytic cleavage with SUMO protease at 4 °C overnight. SEC was performed to separate CtVps391-500 from the expression tag and SUMO protease and yielded >95% pure protein. To obtain phase information for structure determination, selenomethionine-substituted CtVps391-500 was prepared according to well-established methods ^68^.

### Limited proteolysis and protein crystallization

Since initial crystallization approaches did not yield protein crystals suitable for X-ray structure determination, flexible parts of CtVps391-500 were removed by limited proteolysis with α- Chymotrypsin (Merck) at 37° C for 10 minutes. Limited proteolysis was stopped by addition of protease inhibitors (Pierce™ Protease Inhibitor Tablets, EDTA-free, Thermo Fisher Scientific), and an additional SEC in buffer containing 20 mM BIS-TRIS pH 6.5, 200 mM NaCl and protease inhibitors (1:1000) was performed as a final polishing step. Best crystals were obtained by seeding at 12° C and a protein concentration of 7.5 mg/ml in a crystallization condition containing 0.1 M MES pH 7.25 and 20% PEG 2000 MME. Selenium-derivative crystals were flash-cooled in liquid nitrogen in latter condition with 25% glycerol as cryoprotectant.

### Crystal structure determination

Anomalous X-ray data was collected from a single crystal at 100 K at beamline P13, EMBL Hamburg, Germany and diffraction data was processed using XDSAPP3 ^69^. Phase determination using single-wavelength anomalous dispersion at the selenium peak was not successful. A manually trimmed AlphaFold ^61^ model of CtVps39 1-500 was used as a search template in molecular replacement and yielded a single solution with a TFZ score of 24.8 in phenix.phaser ^64,70,71^, which was then used for MR-SAD phasing in phenix.autosol ^64,71,72^ and subsequent density modification and automated model building using phenix.autobuild ^64,71,72^. Iterative cycles of model building in COOT ^63^ and refinement using phenix.refine ^64,72,73^ led to a final model of CtVps391-500 with Rfactors of Rwork 26.9% and Rfree 29.8%. The model contained no Ramachandran outliers with 96.74% residues within favoured regions. Crystallographic statistics are summarized in Table S3. PYMOL and UCSF ChimeraX was used for visualization and graphical analysis.

## Acknowledgments

The authors thank Kristian Parey for his help with model building, and all members of the Ungermann and Moeller lab for feedback. This work was supported by the SFB 944 (P11 to C.U.; P16 to D.K., P20 to F.F., P26 to A.M.), the DFG INST190/196-1 FUGG (A.M.) and the BMBF/DLR 01ED2010 (A.M.).

## Authors‘ contributions

D.S., J.S. and C.K. conceived and designed all experiments together with A.M. and C.U.. D.S. conducted all cryo-EM analyses with support in data analysis and optimization by D.J. and K.S.. J.S. and C.K. did all biochemical experiments with support by A.P.. C.K. did the fusion assays with support by L.L., N.F. did the tethering assays. Protein crystallography and analysis was done by S.K. and D.K. The manuscript was written by D.S., A.M. and C.U. with contributions from all authors.

## Data and materials availability

The electron density maps and models of the yeast HOPS complex and the *C. thermophilum* Vps39 β-propeller have been deposited in the Electron Microscopy Data Bank with ID EMD-XXX and in the PDB with XXX, respectively.

## Notes

### Competing Interest Statement

The authors have declared no competing interest.

## References

1. Ballabio, A. & Bonifacino, J. S. Lysosomes as dynamic regulators of cell and organismal homeostasis. Nat Rev Mol Cell Bio 21, 101–118 (2019).

2. Saftig, P. & Puertollano, R. How Lysosomes Sense, Integrate, and Cope with Stress. Trends Biochem Sci 46, 97–112 (2020).

3. Klionsky, D. J. et al. Autophagy in major human diseases. Embo J 40, e108863 (2021).

4. Sardana, R. & Emr, S. D. Membrane Protein Quality Control Mechanisms in the Endo-Lysosome System. Trends Cell Biol 31, 269–283 (2021).

5. Beek, J. van der, Jonker, C., Welle, R. van der, Liv, N. & Klumperman, J. CORVET, CHEVI and HOPS – multisubunit tethers of the endo-lysosomal system in health and disease. J Cell Sci 132, jcs189134 (2019).

6. Ungermann, C. & Kümmel, D. Structure of membrane tethers and their role in fusion. Traffic 20, 479–490 (2019).

7. Baker, R. W. et al. A direct role for the Sec1/Munc18-family protein Vps33 as a template for SNARE assembly. Science (New York, NY) 349, 1111–1114 (2015).

8. Baker, R. W. & Hughson, F. M. Chaperoning SNARE assembly and disassembly. Nature Reviews Molecular Cell Biology 17, 465–479 (2016).

9. Südhof, T. C. & Rothman, J. E. Membrane fusion: grappling with SNARE and SM proteins. Science (New York, NY) 323, 474–477 (2009).

10. Jahn, R. & Fasshauer, D. Molecular machines governing exocytosis of synaptic vesicles. Nature 490, 201–207 (2012).

11. Wickner, W. & Rizo, J. A cascade of multiple proteins and lipids catalyzes membrane fusion. Molecular Biology of the Cell 28, 707–711 (2017).

12. Kuhlee, A., Raunser, S. & Ungermann, C. Functional homologies in vesicle tethering. Febs Lett 589, 2487–2497 (2015).

13. Spang, A. Membrane Tethering Complexes in the Endosomal System. Frontiers in Cell and Developmental Biology 4, 35 (2016).

14. Mima, J., Hickey, C. M., Xu, H., Jun, Y. & Wickner, W. Reconstituted membrane fusion requires regulatory lipids, SNAREs and synergistic SNARE chaperones. The EMBO Journal 27, 2031–2042 (2008).

15. D’Agostino, M., Risselada, H. J., Lürick, A., Ungermann, C. & Mayer, A. A tethering complex drives the terminal stage of SNARE-dependent membrane fusion. Nature 551, 634–638 (2017).

16. Zick, M. & Wickner, W. Improved reconstitution of yeast vacuole fusion with physiological SNARE concentrations reveals an asymmetric Rab(GTP) requirement. Molecular Biology of the Cell 27, 2590–2597 (2016).

17. Miao, G. et al. ORF3a of the COVID-19 virus SARS-CoV-2 blocks HOPS complex-mediated assembly of the SNARE complex required for autolysosome formation. Dev Cell 56, 427-442.e5 (2021).

18. Carette, J. E. et al. Ebola virus entry requires the cholesterol transporter Niemann-Pick C1. Nature 1–7 (2011) doi:10.1038/nature10348.

19. Welle, R. E. N. van der et al. Neurodegenerative VPS41 variants inhibit HOPS function and mTORC1-dependent TFEB/TFE3 regulation. Embo Mol Med 13, e13258 (2021).

20. Sanderson, L. E. et al. Bi-allelic variants in HOPS complex subunit VPS41 cause cerebellar ataxia and abnormal membrane trafficking. Brain 144, 769–780 (2021).

21. Rieder, S. E. & Emr, S. D. A novel RING finger protein complex essential for a late step in protein transport to the yeast vacuole. Molecular Biology of the Cell 8, 2307–2327 (1997).

22. Hunter, M. R., Scourfield, E. J., Emmott, E. & Graham, S. C. VPS18 recruits VPS41 to the human HOPS complex via a RING–RING interaction. Biochem J 474, 3615–3626 (2017).

23. Segala, G. et al. Vps11 and Vps18 of Vps-C membrane traffic complexes are E3 ubiquitin ligases and fine-tune signalling. Nat Commun 10, 1833 (2019).

24. Lürick, A. et al. Multivalent Rab interactions determine tether-mediated membrane fusion. Mol Biol Cell 28, 322–332 (2017).

25. Bröcker, C. et al. Molecular architecture of the multisubunit homotypic fusion and vacuole protein sorting (HOPS) tethering complex. Proc National Acad Sci 109, 1991–1996 (2012).

26. Ostrowicz, C. W. et al. Defined Subunit Arrangement and Rab Interactions Are Required for Functionality of the HOPS Tethering Complex. Traffic 11, 1334–1346 (2010).

27. Graham, S. C. et al. Structural basis of Vps33A recruitment to the human HOPS complex by Vps16. Proceedings of the National Academy of Sciences 110, 13345–13350 (2013).

28. Chou, H.-T., Dukovski, D., Chambers, M. G., Reinisch, K. M. & Walz, T. CATCHR and HOPS-CORVET tethering complexes share a similar architecture. Nat Struct Mol Biol 23, 761–763 (2016).

29. Peterson, M. R. & Emr, S. D. The class C Vps complex functions at multiple stages of the vacuolar transport pathway. Traffic (Copenhagen, Denmark) 2, 476–486 (2001).

30. Edvardson, S. et al. Hypomyelination and developmental delay associated with VPS11 mutation in Ashkenazi-Jewish patients. J Med Genet 52, 749 (2015).

31. Zhang, J. et al. A Founder Mutation in VPS11 Causes an Autosomal Recessive Leukoencephalopathy Linked to Autophagic Defects. PLoS genetics 12, e1005848–21 (2016).

32. Langemeyer, L., Perz, A., Kümmel, D. & Ungermann, C. A guanine nucleotide exchange factor (GEF) limits Rab GTPase–driven membrane fusion. J Biol Chem 293, 731–739 (2018).

33. Plemel, R. L. et al. Subunit organization and Rab interactions of Vps-C protein complexes that control endolysosomal membrane traffic. Molecular Biology of the Cell 22, 1353–1363 (2011).

34. Füllbrunn, N. et al. Nanoscopic anatomy of dynamic multi-protein complexes at membranes resolved by graphene-induced energy transfer. Elife 10, e62501 (2021).

35. Cabrera, M. et al. Phosphorylation of a membrane curvature–sensing motif switches function of the HOPS subunit Vps41 in membrane tethering. J Cell Biology 191, 845–859 (2010).

36. Auffarth, K., Arlt, H., Lachmann, J., Cabrera, M. & Ungermann, C. Tracking of the dynamic localization of the Rab-specific HOPS subunits reveal their distinct interaction with Ypt7 and vacuoles. Cell Logist 4, 575 e29191 (2014).

37. Zick, M. & Wickner, W. Phosphorylation of the effector complex HOPS by the vacuolar kinase Yck3p confers Rab nucleotide specificity for vacuole docking and fusion. Molecular Biology of the Cell 23, 3429–3437 (2012).

38. Song, H., Orr, A. S., Lee, M., Harner, M. E. & Wickner, W. T. HOPS recognizes each SNARE, assembling ternary trans-complexes for rapid fusion upon engagement with the 4th SNARE. Elife 9, e53559 (2020).

39. Lürick, A. et al. The Habc Domain of the SNARE Vam3 Interacts with the HOPS Tethering Complex to Facilitate Vacuole Fusion*. J Biol Chem 290, 5405–5413 (2015).

40. Song, H., Orr, A., Duan, M., Merz, A. J. & Wickner, W. Sec17/Sec18 act twice, enhancing membrane fusion and then disassembling cis-SNARE complexes. Elife 6, e26646 (2017).

41. Song, H., Torng, T. L., Orr, A. S., Brunger, A. T. & Wickner, W. T. Sec17/Sec18 can support membrane fusion without help from completion of SNARE zippering. Elife 10, e67578 (2021).

42. Orr, A., Song, H. & Wickner, W. Fusion with wild-type SNARE domains is controlled by juxtamembrane domains, transmembrane anchors, and Sec17. Mol Biol Cell mbcE21110583 (2022) doi:10.1091/mbc.e21-11-0583.

43. Zhang, Y. & Hughson, F. M. Chaperoning SNARE Folding and Assembly. Annu Rev Biochem 90, 1–23 (2021).

44. Ha, J. Y. et al. Molecular architecture of the complete COG tethering complex. Nature structural & molecular biology 23, 758–760 (2016).

45. Kant, R. van der et al. Characterization of the Mammalian CORVET and HOPS Complexes and Their Modular Restructuring for Endosome Specificity. The Journal of biological chemistry 290, 30280–30290 (2015).

46. Solinger, J. A., Rashid, H.-O., Prescianotto-Baschong, C. & Spang, A. FERARI is required for Rab11-dependent endocytic recycling. Nat Cell Biol 22, 213–224 (2020).

47. Lai, Y. et al. Molecular Mechanisms of Synaptic Vesicle Priming by Munc13 and Munc18. Neuron 95, 591-607.e10 (2017).

48. Rizo, J. Molecular Mechanisms Underlying Neurotransmitter Release. Annu Rev Biophys 51, (2022).

49. Stepien, K. P. & Rizo, J. Synaptotagmin-1–, Munc18-1–, and Munc13-1–dependent liposome fusion with a few neuronal SNAREs. Proc National Acad Sci 118, e2019314118 (2021).

50. Stepien, K. P., Xu, J., Zhang, X., Bai, X. & Rizo, J. SNARE assembly enlightened by cryo-EM structures of a synaptobrevin-Munc18-1-syntaxin-1 complex. Biorxiv 2022.03.05.483126 (2022) doi:10.1101/2022.03.05.483126.

51. Montoro, A. G. et al. Vps39 Interacts with Tom40 to Establish One of Two Functionally Distinct Vacuole-Mitochondria Contact Sites. Dev Cell 45, 621-636.e7 (2018).

52. Elbaz-Alon, Y. et al. A dynamic interface between vacuoles and mitochondria in yeast. Developmental cell 30, 95–102 (2014).

53. Hönscher, C. et al. Cellular Metabolism Regulates Contact Sites between Vacuoles and Mitochondria. Dev Cell 30, 86–94 (2014).

54. Montoro, A. G. et al. Subunit exchange among endolysosomal tethering complexes is linked to contact site formation at the vacuole. Mol Biol Cell 32, br14 (2021).

55. Janke, C. et al. A versatile toolbox for PCR-based tagging of yeast genes: new fluorescent proteins, more markers and promoter substitution cassettes. Yeast (Chichester, England) 21, 947–962 (2004).

56. Januliene, D. & Moeller, A. Structure and Function of Membrane Proteins. Methods Mol Biology 2302, 153–178 (2021).

57. Punjani, A., Rubinstein, J. L., Fleet, D. J. & Brubaker, M. A. cryoSPARC: algorithms for rapid unsupervised cryo-EM structure determination. Nat Methods 14, 290–296 (2017).

58. Pettersen, E. F. et al. UCSF Chimera—A visualization system for exploratory research and analysis. J Comput Chem 25, 1605–1612 (2004).

59. Pettersen, E. F. et al. UCSF ChimeraX : Structure visualization for researchers, educators, and developers. Protein Sci 30, 70–82 (2020).

60. Varadi, M. et al. AlphaFold Protein Structure Database: massively expanding the structural coverage of protein-sequence space with high-accuracy models. Nucleic Acids Res 50, D439–D444 (2021).

61. Jumper, J. et al. Highly accurate protein structure prediction with AlphaFold. Nature 596, 583–589 (2021).

62. Kidmose, R. T. et al. Namdinator – automatic molecular dynamics flexible fitting of structural models into cryo-EM and crystallography experimental maps. Iucrj 6, 526–531 (2019).

63. Emsley, P., Lohkamp, B., Scott, W. G. & Cowtan, K. Features and development of Coot. Acta Crystallogr Sect D Biological Crystallogr 66, 486–501 (2010).

64. Liebschner, D. et al. Macromolecular structure determination using X-rays, neutrons and electrons: recent developments in Phenix. Acta Crystallogr Sect D 75, 861–877 (2019).

65. Afonine, P. V. et al. Real-space refinement in PHENIX for cryo-EM and crystallography. Acta Crystallogr Sect D Struct Biology 74, 531–544 (2018).

66. Williams, C. J. et al. MolProbity: More and better reference data for improved all-atom structure validation. Protein Sci 27, 293–315 (2018).

67. Zick, M., Stroupe, C., Orr, A., Douville, D. & Wickner, W. T. Membranes linked by trans-SNARE complexes require lipids prone to non-bilayer structure for progression to fusion. eLife 3, e01879–e01879 (2014).

68. Doublié, S. Preparation of selenomethionyl proteins for phase determination. Methods Enzymol 276, 523–30 (1997).

69. Krug, M., Weiss, M. S., Heinemann, U. & Mueller, U. XDSAPP : a graphical user interface for the convenient processing of diffraction data using XDS. J Appl Crystallogr 45, 568–572 (2012).

70. McCoy, A. J. et al. Phaser crystallographic software. J Appl Crystallogr 40, 658–674 (2007).

71. Adams, P. D. et al. PHENIX: a comprehensive Python-based system for macromolecular structure solution. Acta Crystallogr Sect D Biological Crystallogr 66, 213–221 (2010).

72. Terwilliger, T. C. et al. Decision-making in structure solution using Bayesian estimates of map quality: the PHENIX AutoSol wizard. Acta Crystallogr Sect D Biological Crystallogr 65, 582–601 (2009).

73. Afonine, P. V. et al. Towards automated crystallographic structure refinement with phenix.refine. Acta Crystallogr Sect D Biological Crystallogr 68, 352–67 (2012).

74. Langemeyer, L. et al. A conserved and regulated mechanism drives endosomal Rab transition. Elife 9, e56090 (2020).

75. Edgar, R. C. MUSCLE: multiple sequence alignment with high accuracy and high throughput. Nucleic Acids Res 32, 1792–1797 (2004).

76. Waterhouse, A. M., Procter, J. B., Martin, D. M. A., Clamp, M. & Barton, G. J. Jalview Version 2—a multiple sequence alignment editor and analysis workbench. Bioinformatics 25, 1189–1191 (2009).

