## Supplementary Material for "Structure of the lysosomal membrane fusion machinery"

**This PDF file includes:**

Extended Data Figures 1-9

Tables S1-S3

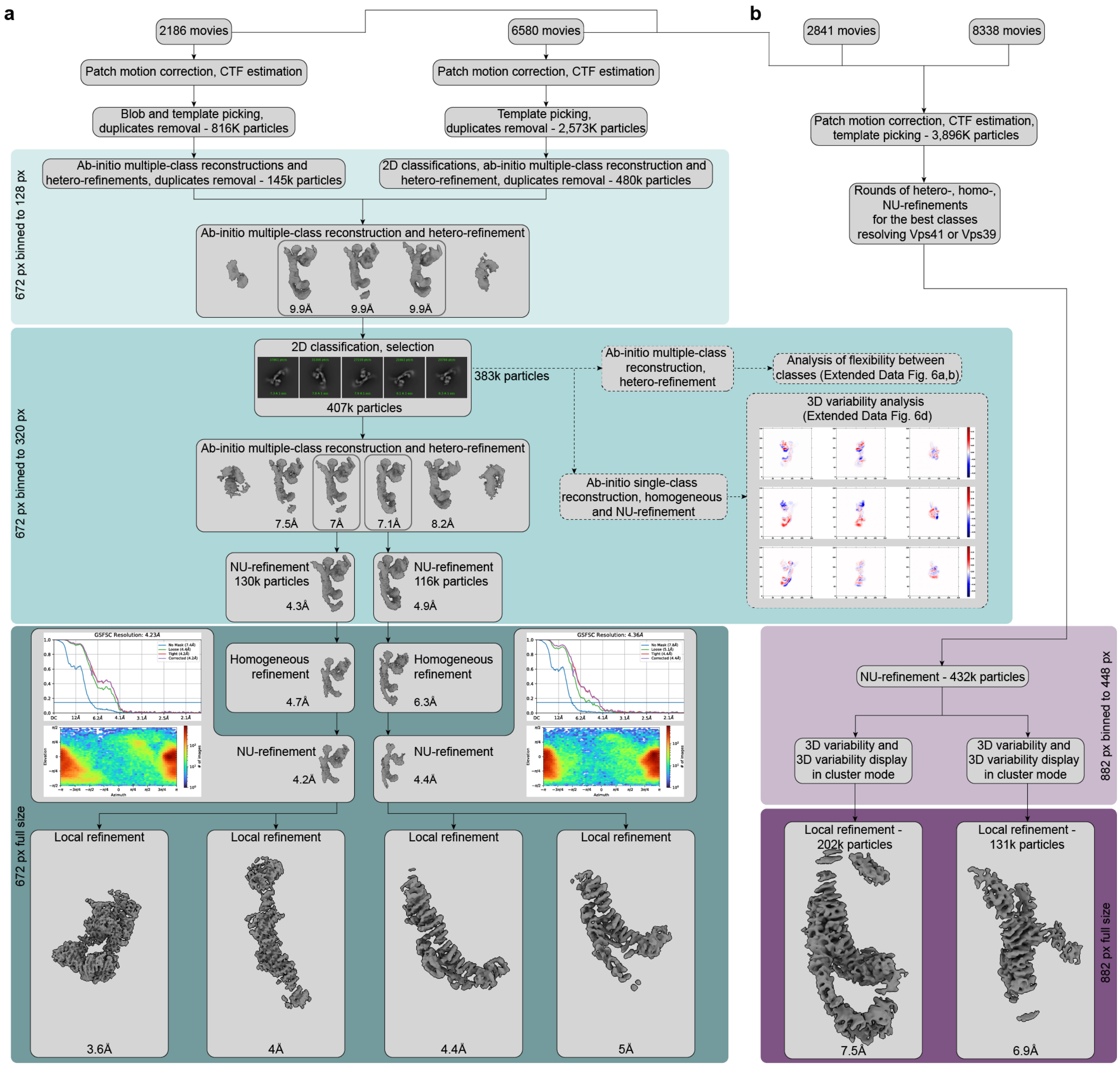


**Extended Data Figure 1: Cryo-EM data processing workflow.**

**a,** Processing of the core part of the complex. 2,186 and 6,580 movies from two different datasets were preprocessed in parallel and used for initial particle sorting by 2D and 3D classifications using the box size of 128 pixels with 4.85 Å per pixel. After initial classifications, approximately 625,000 particles were selected and combined for further processing. A round of ab-initio reconstruction followed by hetero-refinement with five classes was performed to separate the best particles for further refinements. About 407,000 particles from three classes, which reached 9.9 Å resolution (Nyquist for the binned data), were extracted using the box size of 320 pixels with 1.94 Å per pixel, and another round of 2D classification was performed. Afterwards, a new round of ab-initio reconstruction and hetero-refinement with six classes was conducted. Particles from two classes that reached the best resolutions (7 and 7.1 Å) were further processed separately. The first of the selected two classes was refined by NU-refinement to 4.3 Å resolution, while the second class was refined to 4.9 Å. Subsequently, particles were extracted in full box size (672 pixels with 0.924 Å per pixel) and used for further homogeneous and NU-refinement, which resulted in two consensus maps, better resolving either upper or lower part of HOPS (reached 4.2 Å and 4.4 Å resolution, respectively). FSC curves generated in cryoSPARC are shown for both consensus maps. Afterwards, local refinements were applied to improve each of the consensus maps. Local refinements of the first map resulted in two maps covering the SNARE-binding module (3.6 Å resolution) and the upper part for the core of HOPS (4 Å resolution). Local refinements of the second consensus map provided maps of the bottom of HOPS core better covering either Vps18 (4.4 Å resolution) or Vps39 (5 Å resolution). The subset of particles used for generation of both consensus maps was in parallel probed for flexibility using 3D classification (Extended Data Fig. 7a,b) and 3D variability analyses (Extended Data Fig. 7d) (dashed arrows; see Methods for details). **b**, Processing of the distal parts of HOPS. Movies used for core reconstruction (see **a**) were combined with movies from two additional datasets (2841 and 8338 movies, respectively) and preprocessed. After template picking, about 3.9 million particles were selected and subjected to several rounds of heterogeneous, homogeneous and NU-refinements in order to select classes best resolving upper (Vps41) and lower (Vps39) distal parts of HOPS. Afterwards, selected particles were used for a round of NU-refinement with a box size of 448 px with 1.82 Å per pixel. The particles from this NU-refinement were then subjected to 3D variability analysis with masks covering either the Vps41 or Vps39 volume. This was followed by 3D variability display in the cluster mode in order to further select particles. Finally, particles from best clusters were subjected to local refinements at a full box size of 882 pixels with 0.924 Å per pixel resulting in the maps resolving N-terminal regions of Vps39 (7.5 Å resolution) or Vps41 (6.9 Å resolution).

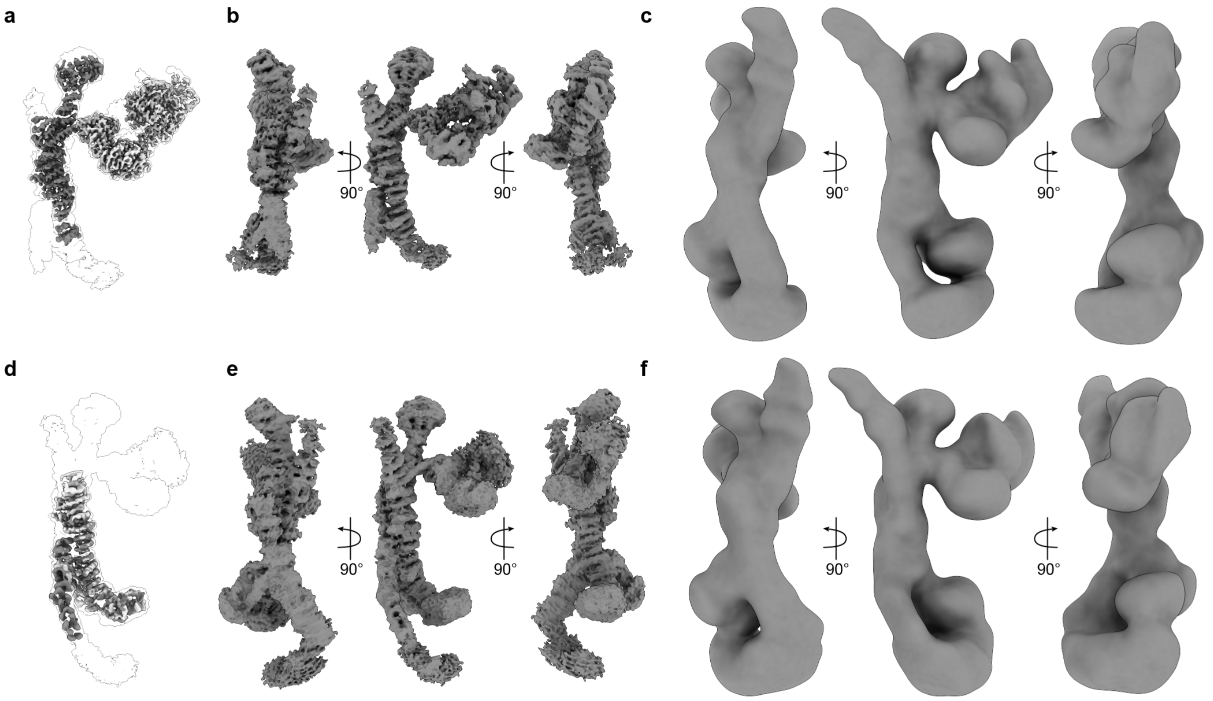


**Extended Data Figure 2: Consensus maps and corresponding local refinement maps.**
**a**, Consensus cryo-EM map of the upper part of HOPS, shown as a transparent envelope, with two corresponding local refinement maps fitted (coloured). **b**, the same consensus map, as in **a**, shown from different sides. **c**, the low-pass-filtered consensus map from **a**, used in Fig. 1e and viewed at a threshold, which allows to demonstrate the densities of N-terminal regions of Vps39 and Vps41. **d**, The consensus map of the lower part of HOPS, shown as a transparent envelope, with two corresponding local refinement maps fitted (coloured). **e**, The consensus map from **d**, shown from different sides. **f**, The map from **d**, shown as in **c**.

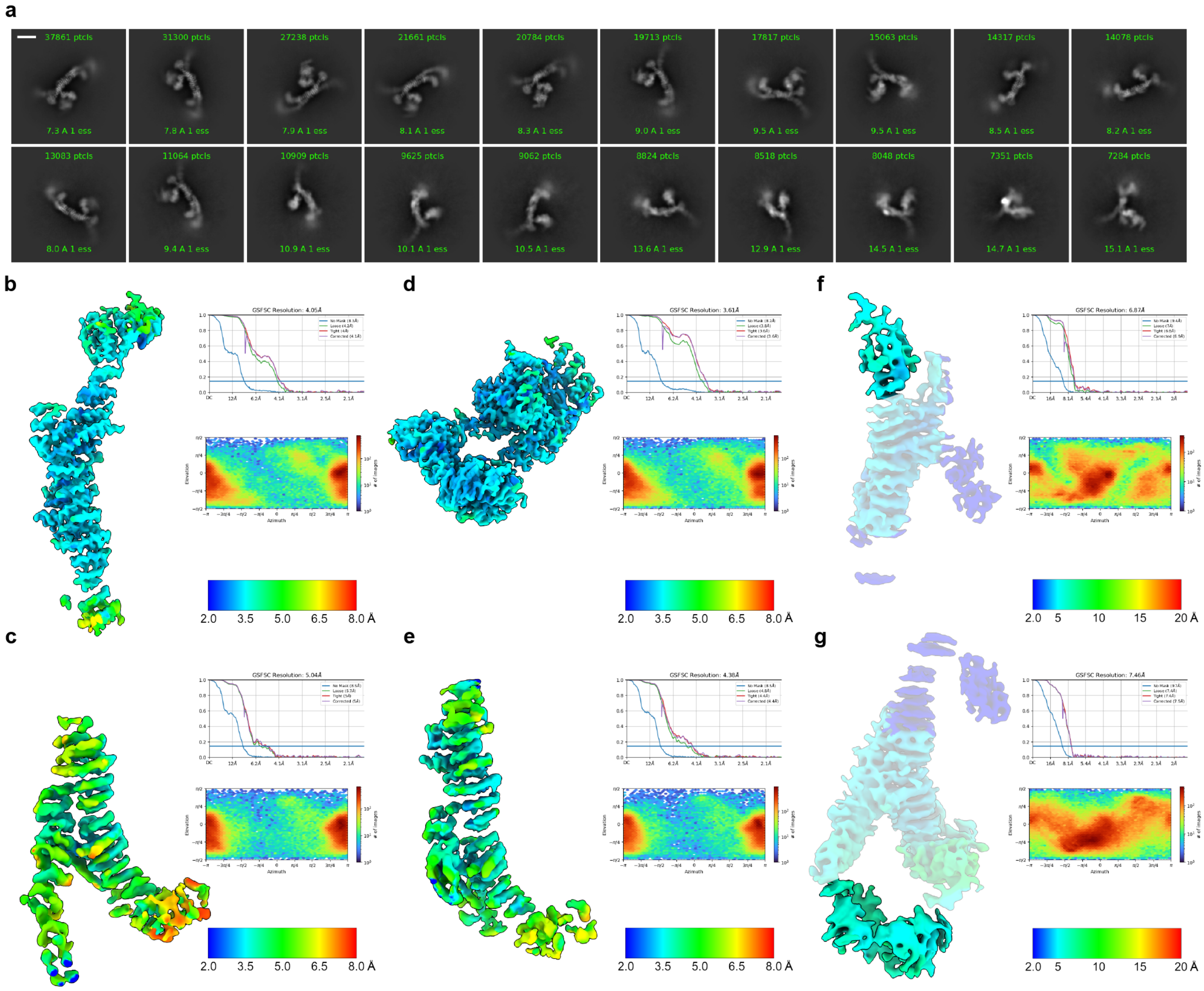


**Extended Data Figure 3: Cryo-EM analysis of HOPS.**

**a**, representative 2D class averages. **b-g**, Local-resolution estimation, FSC curves and angular distribution plots generated in cryoSPARC for each of six local refinement maps (see Extended Data Fig. 1, 2 and Methods). Note, that in **f** and **g**, local resolution was not calculated for the peripheral areas of the maps due to mask limitations and is displayed in dark blue.

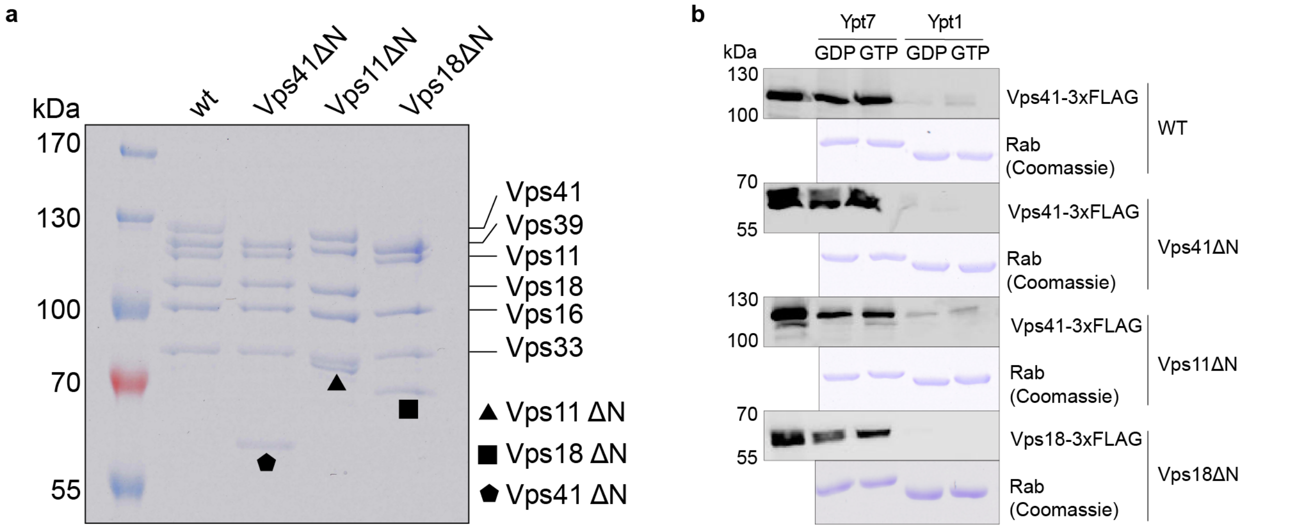


**Extended Data Figure 4: Biochemical analysis of HOPS mutants lacking N-terminal β-propellers.**

**a**, SDS-PAGE of purified wild-type and mutant complexes. **b**, Ypt7-interaction. HOPS wild-type and mutant complexes were added to immobilized GST-Ypt7 or Ypt1 loaded with GTP or GDP. Eluted proteins were analysed by SDS-PAGE and Western blotting. Coomassie gels of corresponding Rab GTPases are shown below.

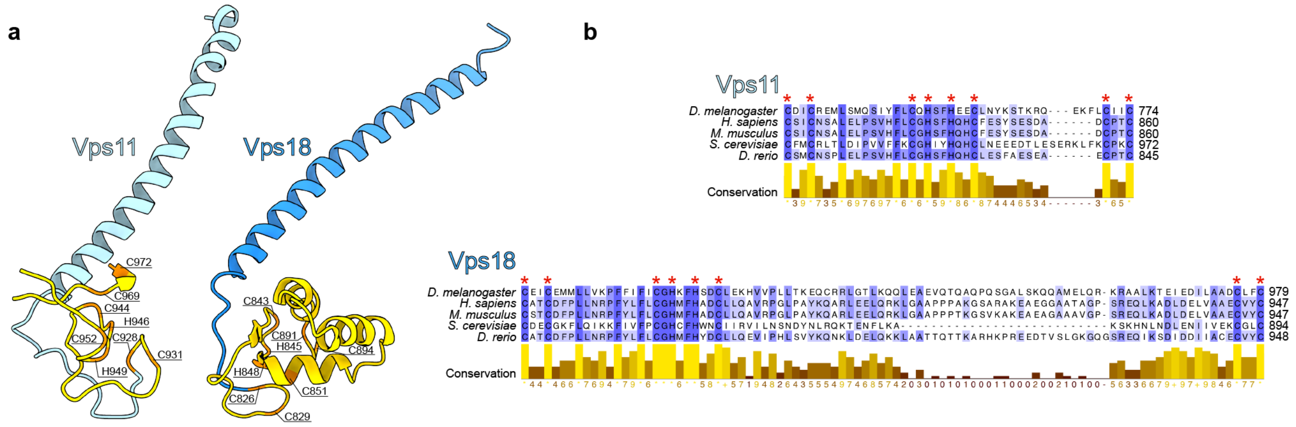


**Extended Data Figure 5: Comparison of Vps11 and Vps18 RING finger domains.**

**a,** Fragments of Vps11 and Vps18 depicting RING finger domains (yellow) following the long helices. Amino acids (orange) participating in folding of the RING finger domains are labeled. **b**, Multiple sequence alignment of Vps11 and Vps18 RING finger domains from different model organisms (*Drosophila melanogaster*, *Homo sapiens*, *Mus musculus*, *Saccharomyces cerevisiae*, *Danio rerio*), made using MUSCLE [75] and visualized in Jalview [76], is shown; colouring by conservation. Conserved amino acids from a are labeled by red asterisks.

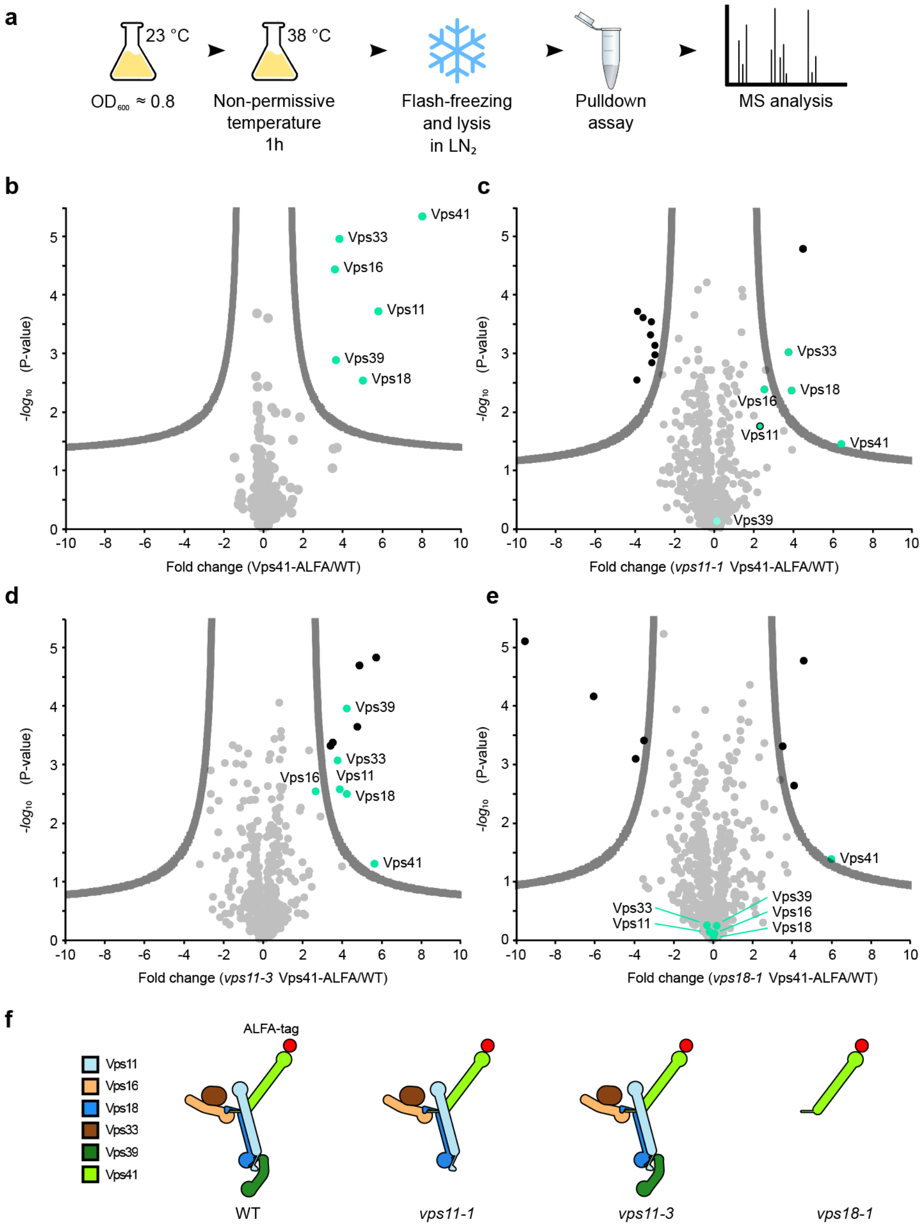


**Extended Data Figure 6: Key role of the RING finger domains of Vps11 and Vps18 in HOPS stability.**

**a,** cartoon of cell growth, lysis, ALFA pull down and mass spectrometry analysis. See Methods for details. **b-e,** Mass spectrometry analysis of Vps41 purified via the ALFA tag from the indicated strains and enriched proteins (green dots). Results of purification from wt (**b**), *vps11-1* (**c**), *vps11-3* (**d**) and *vps18* (**e**) cells. **f**, schematic representation of the results based on the structural model (as in Fig. 1g).

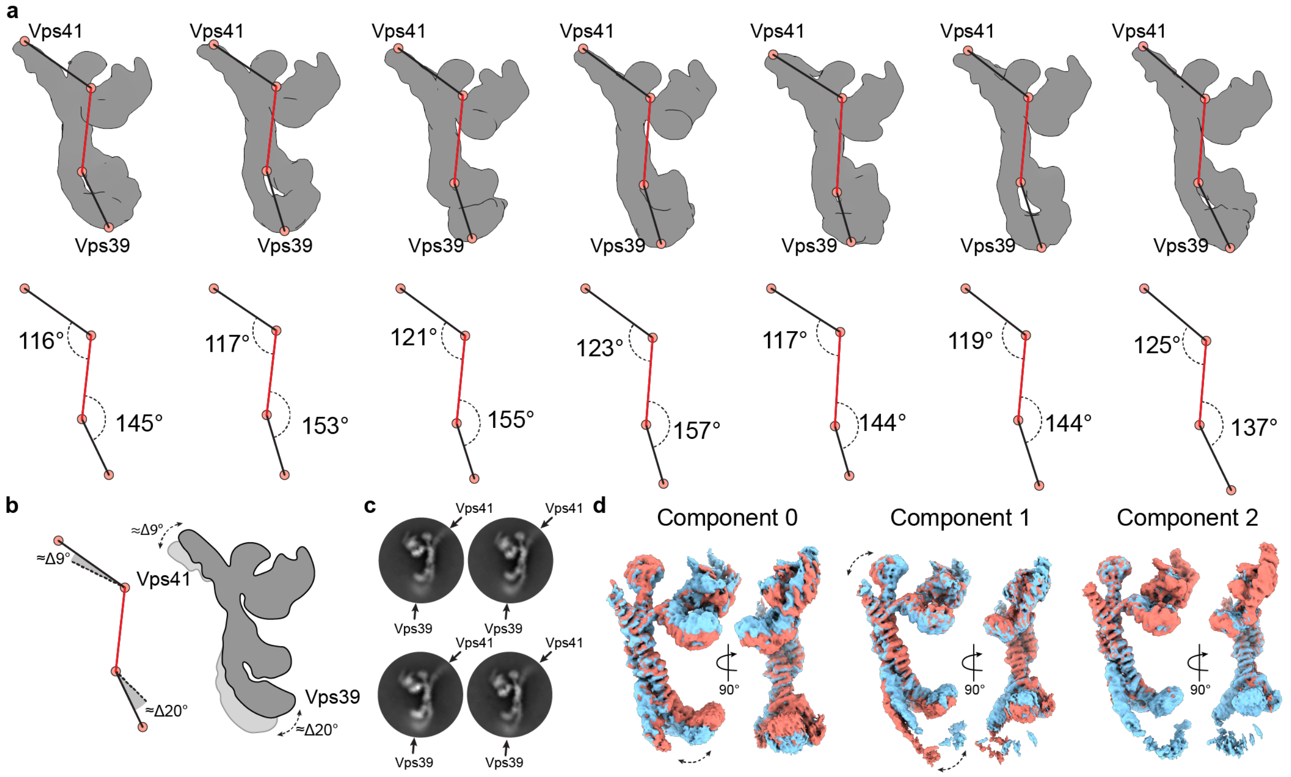


**Extended Data Figure 7: Flexibility analysis of the HOPS complex.**

**a**, upper row, maps generated by a hetero-refinement with multiple classes from particles used for local refinements (see Extended Data Fig. 1a). All classes demonstrate different positions of densities at the distal ends relative to the rigid core (highlighted by red lines) of the volumes. Lower row, rotation angles calculated between the points indicated on the maps from above. **b**, Angular difference calculated from angles in **a**, demonstrating limited movements of Vps41 and Vps39, as depicted in the cartoon to the right. **c**, representative 2D class averages show fuzzy densities of Vps41 and Vps39 owing to their flexibility. **d**, 3D variability analysis of particles used for local refinements (see Extended Data Fig. 1a). 3D density maps at negative (red) and positive (blue) positions along each variability component are shown.

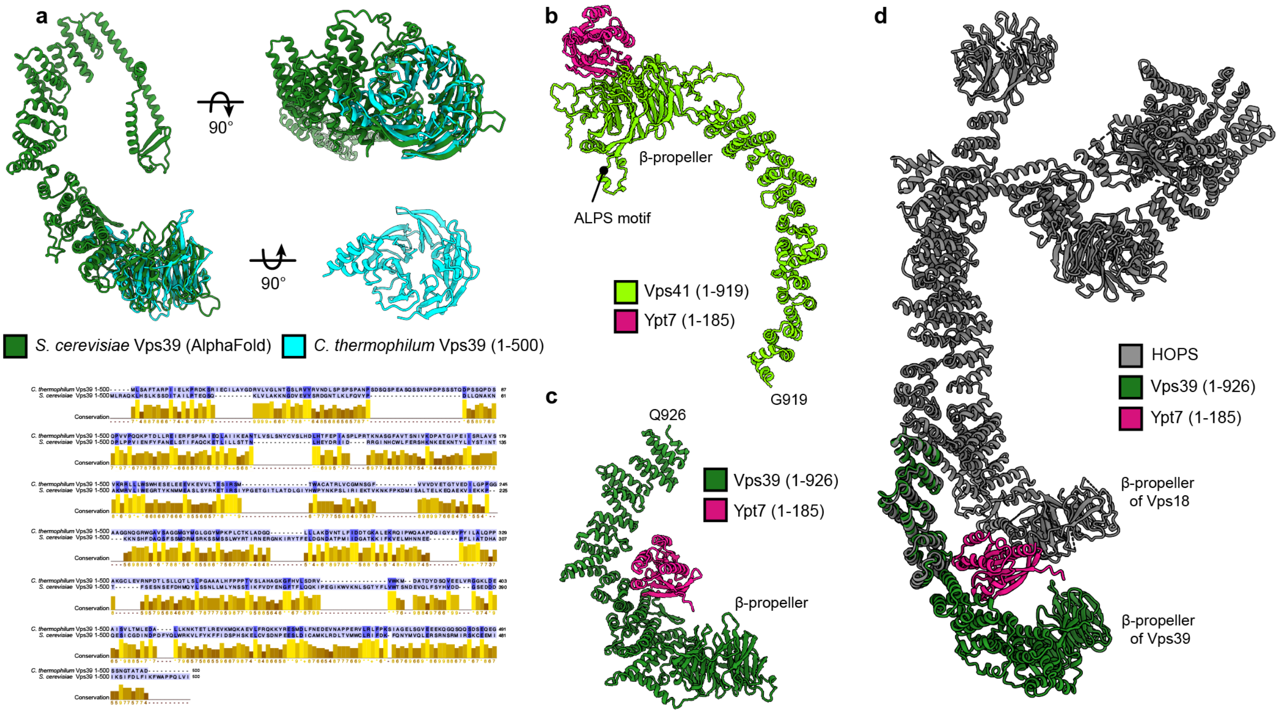


**Extended Data Figure 8: Ypt7 interaction with Vps41 and Vps39.**

**a**, Structure of the β-propeller of *C. thermophilum* Vps39 confirms the structure prediction. Cristal structure of *C. thermophilum* Vps39 (1-500) construct (cyan) is shown and superimposed on the model of *S. cerevisiae* Vps39 (dark green) generated by AlphaFold. Below, sequence alignment of Vps39 N-terminal fragments from *C. thermophilum* and *S. cerevisiae* is displayed. Alignment is made using MUSCLE [75] and visualized in Jalview [76]; colouring by conservation. **b**, Model of Vps41 (light green, residues 1-919) with Ypt7 (pink, residues 1-185) generated by AlphaFold (multimer mode). According to the model, Ypt7 binds to the β-propeller of Vps41 at the opposite side than the identified membrane-interacting ALPS motif [35]. **c**, Model of Vps39 (dark green, residues 1-926) with Ypt7 (pink, residues 1-185) was obtained as in **a**. Ypt7 binds to the inside of the α-solenoid of Vps39 according to the model. **d**, The model from **b**, fitted to the atomic model of HOPS generated in this study (gray, see Fig. 2). According to the AlphaFold model, Ypt7 has a contact site not only with Vps39 but also with Vps18 β-propeller.

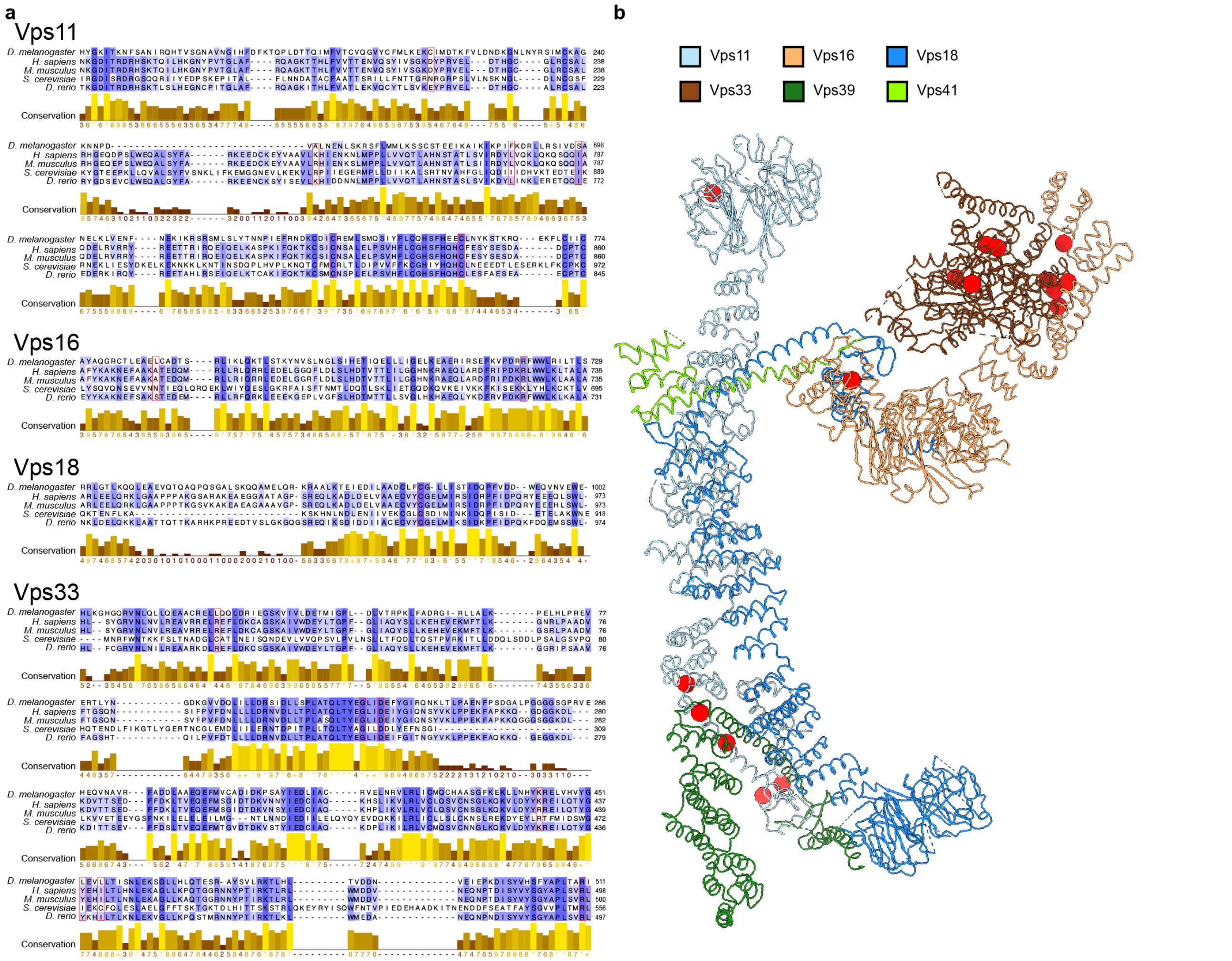


**Extended Data Figure 9: Point mutations affecting HOPS function.**

**a,** Multiple sequence alignment of HOPS subunit fragments from different model organisms (*Drosophila melanogaster*, *Homo sapiens*, *Mus musculus*, *Saccharomyces cerevisiae*, *Danio rerio*) displaying known point mutations (red frames) [5,29]. **b**, Mutations from **a** are labeled on the structure of HOPS (see Fig. 2) as red spheres. In cases where mutations were described for a different organism, respective homologous residues in yeast were mapped according to the alignment in **a**.

**Table S1. Yeast strains used in the study**

| Strain | Genotype | Source |
|---|---|---|
| SEY6210 | MATalpha *leu2-3,112 ura3-52 his3-Δ200 trp-Δ901 lys2-801 suc2-Δ9 GAL* | Reggiori Lab |
| CUY2489 | MATalpha *his3-Δ200 leu2-Δ0 lys2-Δ0 met15-Δ0 trp1-Δ63 ura3-Δ0 VPS11::HIS3-GAL1pr VPS16::NatNT2-GAL1pr VPS18::KanMX-GAL1pr* | Ostrowicz et al., 2010 |
| CUY13050 | MATa *his3-Δ200 met15-Δ0 trp1-Δ63 ura3-Δ0 VPS41::TRP1-GAL1pr VPS39::KanMX-GAL1pr VPS33::HIS3-Gal1pr VPS41::HphNT1-FLAG* | This study |
| CUY13080 | CUY2489xCUY13050 | This study |
| CUY13079 | MATa *his3-Δ200 leu2-Δ0 met15-Δ0 trp1-Δ63 ura3-Δ0 VPS39::KanMX- Gal1pr VPS33::HIS3-GAL1pr VPS41Δaa1-499::CloNAT- GAL1pr VPS41ΔN::3xFLAG-HphNT1* | This study |
| CUY13194 | MATalpha *his3-Δ200 leu2-Δ0 lys2-Δ0 met15-Δ0 trp1-Δ63 ura3-Δ0 VPS11::HIS3-GAL1pr VPS16::NatNT2-GAL1pr VPS18::KanMX-GAL1pr-3HA VPS41::Δaa1-499-TRP1* | This study |
| CUY13195 | CUY13079xCUY13194 | This study |
| CUY13751 | MATalpha *his3-Δ200 leu2-Δ0 lys2-Δ0 met15-Δ0 trp1-Δ63 ura3-Δ0 VPS11::HIS3-GAL1pr VPS18::TRP-GAL1pr VPS39::KanMX-GAL1pr VPS39::FLAG-HphNT1* | This study |
| CUY13764 | Matalpha *his3-Δ200 leu2-Δ0 lys2-Δ0 met15-Δ0 trp1-Δ63 ura3-Δ0 VPS11Δaa1-349::URA-GAL1pr VPS16::NatNT2-GAL1pr VPS18::KanMX-GAL1pr-3HA VPS39::TRP1-GAL1pr VPS39::KanMX-GAL1pr VPS33::HIS3-Gal1pr VPS41::HphNT1-FLAG* | This study |
| CUY1908 | MATa *his3-Δ200 met15-Δ0 trp1-Δ63 ura3-Δ0 VPS392::TRP1-GAL1pr VPS41::KanMX-GAL1pr Vps33::HIS3-Gal1pr* | Ungermann lab |

| | | |
|---|---|---|
| CUY13751 | MATalpha *his3-Δ200 leu2-Δ0 lys2-Δ0 met15-Δ0 trp1-Δ63 ura3-Δ0 VPS11::HIS3-GAL1pr VPS16::kanMX-GALpr1 VPS18Δaa1-335::TRP-GAL1pr VPS18::FLAG-HphNT1* | This study |
| CUY13763 | CUY1908xCUY13763 | This study |
| CUY2742 | Mat alpha *vps11-1(Ts) leu2-3,112 ura3-52 his3-Δ200 trp1-Δ901 lys2-801 suc2-Δ9 VPS11-1::HIS3-HA* | Robinson et al. 1991 |
| CUY2743 | Mat alpha *vps11-3(Ts) leu2-3,112 ura3-52 his3-Δ200 trp1-Δ901 lys2-801 suc2-Δ9* | Robinson et al. 1991 |
| CUY2744 | Mat alpha *vps18-1(Ts) leu2-3,112 his3-Δ200 trp1-Δ901 lys2-801 suc2-Δ9* | Robinson et al. 1991 |
| CUY13557 | MATalpha *leu2-3,112 ura3-52 his3-Δ200 trp-Δ901 lys2-801 suc2-Δ9 VPS41::ALFA-HphNT1* | This study |
| CUY13598 | Mat alpha *leu2-3,112 ura3-52 his3-Δ200 trp1-Δ901 lys2-801 suc2-Δ9 VPS11-1::HIS3-HA VPS41::ALFA-HphNT1* | This study |
| CUY13559 | Mat alpha *leu2-3,112 ura3-52 his3-Δ200 trp1-Δ901 lys2-801 suc2-Δ9 VPS41::ALFA-HphNT1* | This study |
| CUY13561 | Mat alpha *leu2-3,112 his3-Δ200 trp1-Δ901 lys2-801 suc2-Δ9 VPS41::ALFA-HphNT1* | This study |

References

1) Ostrowicz, C. W. *et al.* Defined Subunit Arrangement and Rab Interactions Are Required for Functionality of the HOPS Tethering Complex. *Traffic* 11, 1334–1346 (2010).
2) Robinson, J.S. et al. A putative zinc finger protein, Saccharomyces cerevisiae Vps18p, affects late Golgi functions required for vacuolar protein sorting and efficient alpha-factor prohormone maturation. *Mol Cell Biol* 11(12), 5813-5824 (1991).

### Table S2. Cryo-EM data collection, refinement and validation statistics

| | HOPS upper<br>(EMDB-xxxx: consensus upper,<br>EMDB-xxxx: SNARE-binding,<br>EMDB-xxxx: backbone)<br>(PDB xxxx) | HOPS lower<br>(EMDB-xxxx: consensus lower,<br>EMDB-xxxx: bottom-Vps18,<br>EMDB-xxxx: bottom-Vps39)<br>(PDB xxxx) |
|---|---|---|
| **Data collection and processing** | | |
| Magnification | 130,000 | 130,000 |
| Voltage (kV) | 200 | 200 |
| Electron exposure (e–/Å$^2$) | 50 | 50 |
| Defocus range (μm) | -0.8 to -2.8 | -0.8 to -2.8 |
| Pixel size (Å) | 0.924 | 0.924 |
| Symmetry imposed | C1 | C1 |
| Initial particle images (no.) | 2,565,533 (after duplicates removal) | 2,565,533 (after duplicates removal) |
| Final particle images (no.) | 129,388 (129,389 for consensus upper) | 115,273 (115,272 for bottom-Vps39) |
| Map resolution (Å) | 4.2 (consensus upper), 3.6 (SNARE-binding), 4.0 (backbone) | 4.4 (consensus lower), 4.4 (bottom-Vps18), 5.0 (bottom-Vps39) |
| FSC threshold | 0.143 | 0.143 |
| **Refinement** | | |
| Initial model used | AlphaFold | AlphaFold |
| Model resolution range (Å) | 3.6-5.0 | 3.6-5.0 |
| FSC threshold | 0.143 | 0.143 |
| Map sharpening *B* factor (Å$^2$) | -90.6 (consensus upper)<br>-76.3 (SNARE-binding)<br>-78.7 (backbone) | -111.4 (consensus lower)<br>-116.1 (bottom-Vps18)<br>-198.2 (bottom-Vps39) |
| Model composition | | |
| Non-hydrogen atoms | 25801 | 25801 |
| Protein residues | 3581 | 3581 |
| R.m.s. deviations | | |
| Bond lengths (Å) | 0.005 | 0.005 |
| Bond angles (°) | 0.979 | 0.979 |
| Validation | | |
| MolProbity score | 2.08 | 2.08 |
| Clashscore | 11.78 | 11.78 |
| Poor rotamers (%) | 0 | 0 |
| Ramachandran plot | | |
| Favored (%) | 91.87 | 91.87 |
| Outliers (%) | 0 | 0 |

**Table S3. Crystallographic data collection and refinement statistics (molecular replacement)**

| | ***Ct*Vps39$_{1-500}$ (SAD)** (pdb code: XXX) |
|---|---|
| **Data collection** | |
| X-ray source | EMBL, Hamburg, Germany, P13 |
| Wavelength (Å) | 0.97937 |
| Space group | P21212 |
| Cell dimensions | |
| *a*, *b*, *c* (Å) | 64.70, 102.75, 59.56 |
| Resolution (Å) | 43.82-2.89 (3.07-2.89)* |
| Total reflections | 114239 (16970) |
| Multiplicity | 6.72 (6.48) |
| Unique reflections | 17010 (2619) |
| Completeness (%) | 99.1 (94.8) |
| $R_{meas}$ (%) | 23.1 (124.3) |
| CC(1/2) | 99.3 (76.6) |
| $I / \sigma I$ | 7.70 (1.80) |
| Mosaicity (°) | 0.523 |
| Wilson *B*-factor (Å$^2$) | 58.48 |
| **Refinement** | |
| Resolution (Å) | 40.23-2.89 |
| Reflections (working, test set) | 9264, 463 |
| $R_{factor}$, $R_{free}$ | 0.269, 0.298 |
| Completeness for range (%) | 99.1 |
| No. atoms | |
| Protein | 2681 |
| Solvent | 10 |
| Protein residues | 345 |
| R.m.s. deviations | |
| Bond lengths (Å) | 0.003 |
| Bond angles (°) | 0.551 |
| mean *B* value (Å$^2$) | 58.1 |

*Values in parentheses are for highest-resolution shell.
